# A new mathematical model of phyllotaxis to solve the genuine puzzle spiromonostichy

**DOI:** 10.1101/2021.09.03.458814

**Authors:** Takaaki Yonekura, Munetaka Sugiyama

## Abstract

The view is widely accepted that the inhibitory effect of existing leaf primordia on new primordium formation determines phyllotactic patterning. Previous studies have shown that mathematical models based on such inhibitory effect can generate most of phyllotactic patterns. However, a few types of phyllotaxis still remain unaddressed. A notable example is costoid phyllotaxis showing spiromonostichy, which is characterized by a steep spiral with a small divergence angle and is unique to Costaceae plants. Costoid phyllotaxis has been called a “genuine puzzle” because it seems to disagree with the inhibitory effect-based mechanism. In an attempt to produce a steep spiral pattern, we developed a new mathematical model assuming that each leaf primordium emits not only the inhibitory effect but also some inductive effect. Computer simulations with the new model successfully generated a steep spiral pattern when these two effects met a certain relationship. The obtained steep spiral matched the real costoid phyllotaxis observed with *Costus megalobractea*. We also found by the mathematical model analysis that the early phyllotactic transition in the seedlings of this plant can be explained by the SAM enlargement.

## Introduction

The arrangement of leaves around the stem, termed phyllotaxis, is one of the most conspicuous patterns seen in the plant architecture because of its geometric beauty. There are various types of phyllotaxis but the variety is rather limited. Most plant species show one of the four common types of phyllotaxis: distichous, Fibonacci spiral, decussate, and tricussate (Yonekura and Sugiyama, 2021), where Fibonacci spiral represents spiral phyllotaxis with the divergence angle close to the golden angle, 137.5°, and with a mathematical relation to Fibonacci sequence (Jean, 1994). The limited variation suggests some constraints coming from the regulatory mechanism behind phyllotactic pattern formation. In this respect, as early as one and a half centuries ago, Hofmeister described an empirical law of leaf primordium positioning (Hofmeister, 1868) and is now known as Hofmeister’s axiom (Jean, 1994). It states that, on the periphery of the shoot apical meristem (SAM), “each leaf arises in the largest gap between existing leaves or primordia and as far away as possible from them” (Jean, 1994).

Following Hofmeister’s axiom, various models assuming repulsive interaction between leaf primordia were proposed to explain phyllotactic pattern formation. In early models, the repulsive effect of each primordium was represented by a solid disk surrounding it, which was not allowed to overlap with the other disk (Snow and Snow, 1962; Adler, 1974; Mitchison 1977; Roberts 1977). The repulsive effect was expressed as a kind of inhibitory field in more generalized models (Mitchison, 1977; Levitov, 1991; Douady and Couder, 1992; 1996a; 1996b). Among these models, two models of Douady and Couder (1996a; 1996b;), referred to as DC1 and DC2 hereafter, were most extensively analyzed and have served as the basic framework for later theoretical studies of phyllotactic pattern formation. Both DC1 and DC2 postulate a constant inhibitory field emanated from the center of each leaf primordium. In DC1, a new primordium arises one by one where the inhibitory field strength is minimal on the SAM periphery at a certain given plastochron, and thus DC1 is specialized for regular patterns of alternate phyllotaxis. In DC2, primordium initiation occurs whenever and wherever the inhibitory field strength falls below a given threshold on the SAM periphery, and thus DC2 can deal with both alternate and whorled patterns. Computer simulations with DC1 demonstrated that specific divergence angles such as the golden angle and 180°are more stable in alternate phyllotaxis, and computer simulations with DC2 successfully produced all major types of phyllotaxis.

In the 2000s, physiological molecular biological studies indicated the critical importance of the plant hormone auxin and its polar transport driven by the auxin efflux carrier PIN1 in the initiation of shoot lateral organs (Reinhardt et al., 2000; 2003; Benková et al., 2003) Further analysis suggested that a positive feedback loop between auxin distribution and PIN1 localization works for the spontaneous generation of auxin convergence in the epidermal layer of the shoot apex and these findings were integrated into mathematical models (Smith et al., 2006, Jönsson et al., 2006). It was also demonstrated by computer simulations that these auxin-based models are able to create major phyllotactic patterns. In these models, the auxin convergence determines the site of primordium initiation and auxin depletion into the convergence from its surroundings by the positive feedback dynamics generates the inhibitory field. The parameters of the auxin-based models were mapped to the parameters of the DC2, which showed that DC2 can be treated as an abstract model of phyllotactic pattern formation regulated by auxin (Mirabet et al., 2012). DC models and auxin-based models provided a basis for understanding the formation of major phyllotactic patterns, they did not address several minor types of phyllotaxis such as orixate phyllotaxis, a tetrastichous alternate type with a four-cycle change of the divergence angle in the sequence of 180°, 90°, 180°, and 270°. Additionally, these models did not fully explain for the overwhelming dominance of Fibonacci spiral in spiral phyllotaxis. In an attempt to address these problems, we constructed expanded versions of DC1 and DC2, which we call EDC1 and EDC2, respectively, by introducing a primordial age-dependent increase of the inhibitory power emission (Yonekura et al., 2019). As a result of computer simulation analysis with EDC models, EDC2 was shown to reproduce broader types of phyllotaxis including orixate phyllotaxis and fit better to the dominance of Fibonacci spiral than the previous models (Yonekura et al. 2019).

Now almost all phyllotactic patterns that occur in nature can be produced by mathematical models, EDC2 in particular. However, there still remain a few exceptions. Of these, the most striking one is costoid phyllotaxis uniquely found in Costaceae, Zingiberales (Fig 1). Costoid phyllotaxis is characterized by spiromonostichy, that is a steep spiral with a small divergence angle (e.g., 47.7° in *Costus spicatus*; Snow, 1952) in which all leaves are lined up in a single oblique row, i.e., monostichy. The spiromonostichous appearance of costoid phyllotaxis clearly distinguishes it from common spiral phyllotaxis. It is also notable that the divergence angle of costoid phyllotaxis is continuously variable during development and between species and is not converged to specific values linked to Fibonacci or related sequences (Kirchoff and Rutishauser, 1990). The small divergence angle of costoid phyllotaxis apparently results from a peculiar spatial relationship between leaf primordia such that a new leaf primordium is positioned near its preceding primordium. Because this feature violates Hofmeister’s axiom, costoid phyllotaxis or spriomonostichy has been called a “famous and fascinating puzzle” (Kirchoff and Rutishauser, 1990) and a “genuine puzzle” (Jean, 1994).

**Fig 1.**
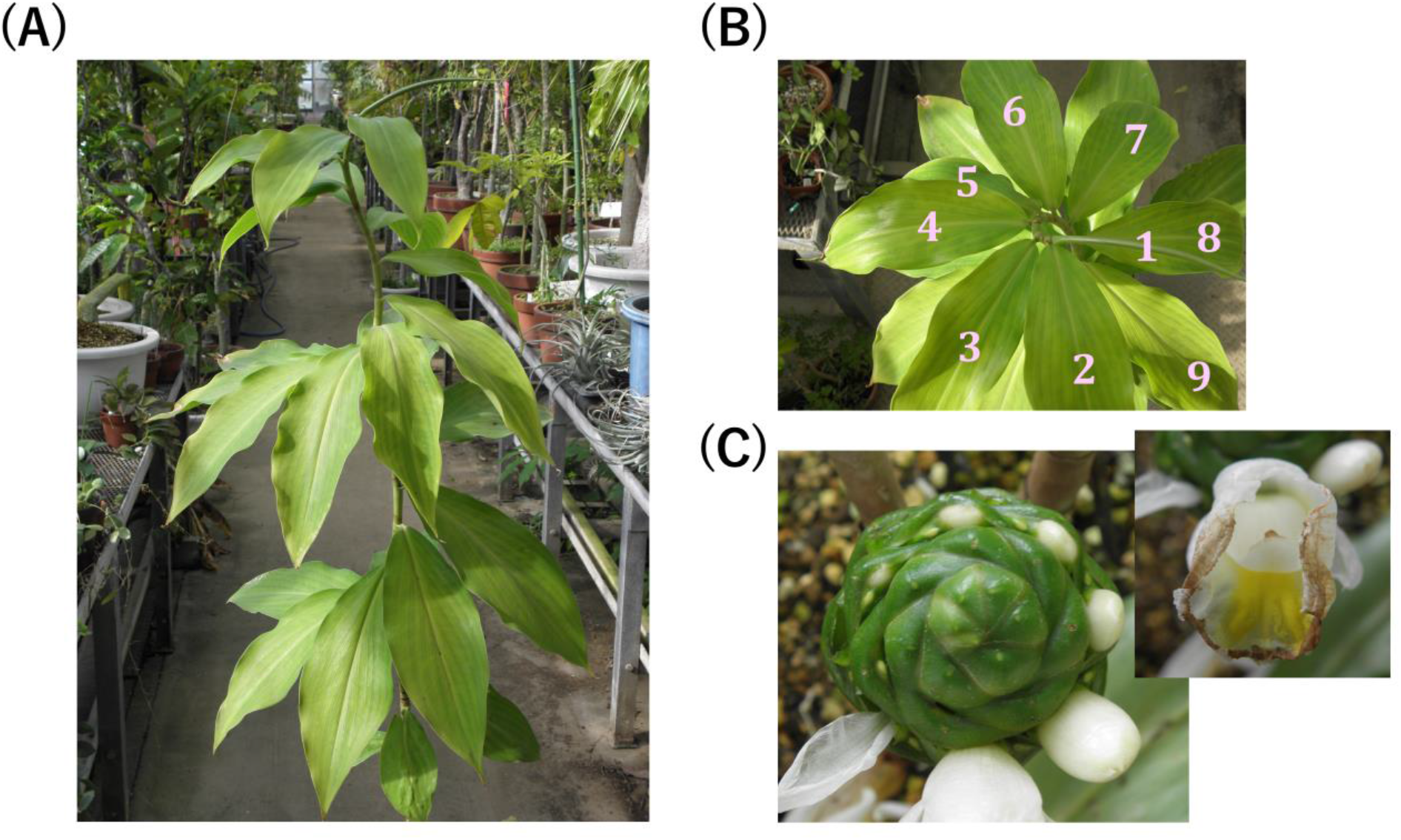
Photographs of *C. megalobractea*. (A) Side view of the shoot of an adult vegetative plant. (B) Top view of the shoot of an adult vegetative plant. (C) Inflorescence and flowers.

In attempts to explain the unusual leaf positioning of costoid phyllotaxis, several ideas have been suggested (Kirchoff and Rutishauser, 1990), such as those assuming a possible effect of the asymmetric young leaf sheath (Schumann, 1892), a possible effect of the pressure from the completely sheathing leaves (Weisse, 1932), a possible loss of one ortho/parastichy from (spiro-)distichous phyllotaxis (Hirmer, 1922; Kirchoff and Rutishauser, 1990), possible congenital torsion of the shoot apex (Goebel 1928; von Veh, 1931; Troll 1937), a possible effect of the presumed prohibition of the simultaneous existence of plural primordia in the apex (Smith, 1941), and a possible effect of the tilt of the shoot apex (Snow, 1952). All of these, however, are just intuitive ideas without having been tested either theoretically or experimentally for their validities, and even some of them obviously contradict morphological observations of costoid phyllotaxis (Kirchoff and Rutishauser, 1990). Jean (1988) proposed a model for systematically relating various phyllotactic patterns including the costoid pattern, but this is an interpretative model and does not give any insights into the mechanism of phyllotactic patterning. Thus, so far, there have been no mechanistic models addressing the generation of the costoid pattern.

In the present study, we analyzed costoid phyllotaxis of *Costus megalobractea* morphologically, examined the ability of the previous models to produce costoid phyllotaxis, and then constructed a new mathematical model by introducing a hypothetical inductive field that surrounds the existing leaf primordia and acts positively for new primordium initiation. The results obtained indicate that a balance between expansions of the induction and inhibition ranges is critical for the generation of costoid phyllotaxis. Computer simulation with our new model under appropriate parameter settings produced realistic costoid phyllotaxis.

## Materials, methods and models

### Plant materials and growth condition

Shoot apices of *C. megalobractea* that had been collected from adult vegetative plants growing in the greenhouse of Koishikawa Botanical Gardens, Graduate School of Science, The University of Tokyo were subjected to morphological analysis of phyllotaxis. For measurement of early changes in the divergence angle, young seedlings of *C. megalobractea* cultured at 25°C under continuous light from white fluorescent lamps were used instead of adult plants.

### Microscopic observation

The shoot apices of *C. megalobractea* were fixed with FAA (5% v/v formalin, 5% v/v acetic acid, 50% v/v ethanol), dehydrated in an ethanol series, and finally infiltrated in 100% ethanol. Then they were embedded in Technovit® 7100, cut into 5-μm-thick sections with a rotary microtome, and stained with 0.5% w/v toluidine blue / 0.1% Sodium carbonate. Images of the sections were assembled using MosaicJ plugin (https://imagej.net/MosaicJ) of ImageJ 1.49v (https://imagej.nih.gov/). For each leaf (primordium), the gravity center of the midvein if the midvein was obvious, or the gravity center of the whole leaf (primordium) section if otherwise was determined and used as its position for morphometric analysis.

### Computer simulations

Model simulations were implemented in C++ with Visual C++ in Microsoft Visual Studio® 2019 as an integrated development environment. Contour mapping was performed with OpenCV ver. 4.1.0 (https://opencv.org/).

Simulation analysis with EDC1 and EDC2 was performed using the codes previously published (Yonekura et al., 2019). The details of these models are described in Text S1. Computer simulations were initiated by placing a single primordium on the SAM periphery and performed with the spatial resolution of 0.1°. All models were simulated with the time step of Δ*t_sim_* = 0.001.

As a distance between points 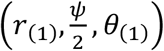 and 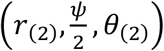 (expressed in spherical coordinate) on the conical surface, instead of the true Euclidian distance, its slightly modified version defined in the following equation is used to avoid the discontinuity problem (Douady and Couder, 1996b).

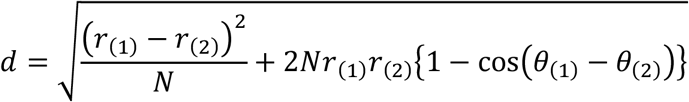

In all model simulations, calculation was iterated while the total number of primordia was less than 100, the standardized plastochron was less than 5, and the passing time in the simulation was less than twice the number of primordia. Alternate and whorled patterns generated by simulation were judged for the stability and regularity of divergence angles and for the number of primordia per node, respectively. Then the patterns were categorized and displayed as shown in Fig 2.

**Fig 2.**
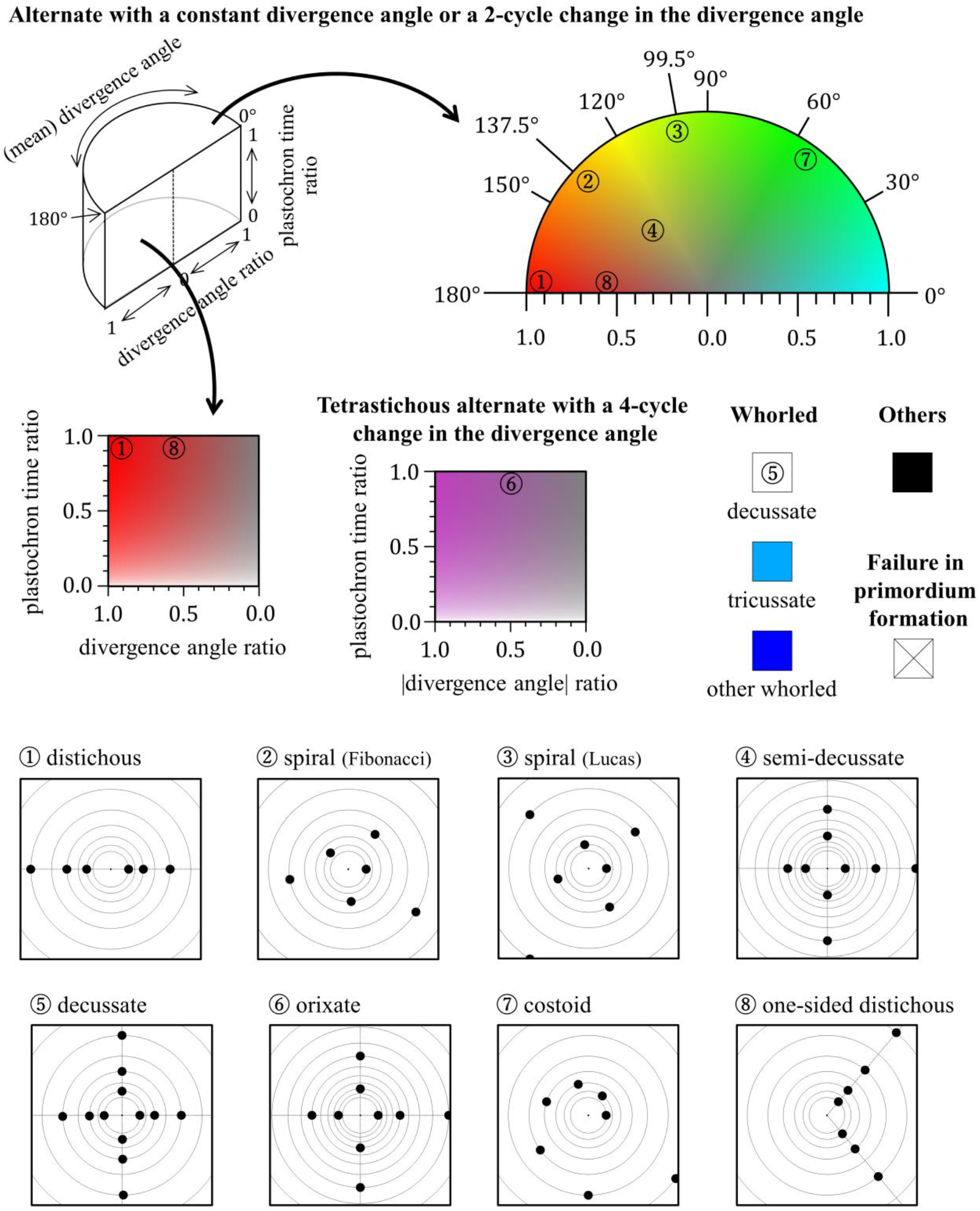
Color legend for the phyllotactic patterns generated in computer simulations. The phyllotactic patterns generated in computer simulations were classified into an alternate pattern with a constant divergence angle or a two-cycle change in the divergence angle; a tetrastichous alternate pattern with a four-cycle change in the divergence angle; a whorled pattern; and other patterns. When no new primordia were formed in simulation, it was indicated by the saltire mark (×). Whorled patterns were further classified into decussate, tricussate, and other whorled patterns. These patterns were distinguished using different colors. For regular alternate patterns with a constant divergence angle, the divergence angle was indicated by a color hue from cyan (0°) to red (180°). In the case of alternate patterns with a two-cycle divergence angle change with a constant absolute value of the divergence angle, the color hue was assigned for the absolute value of the divergence angle. In the case of other alternate patterns with a two-cycle divergence angle change, the color hue was assigned for the mean absolute value of the successive divergence angles. In these two-cycle alternate patterns, small-to-large ratios of two successive plastochron times and two successive divergence angles were represented by lightness (full lightness for 0) and saturation (full saturation for 1), respectively. Tetrastichous alternate patterns with a four-cycle divergence angle change were similarly expressed by color brightness and saturation based on their ratios of plastochron times and divergence angles; however, instead of the divergence angles themselves, the absolute values of divergence angles were used to calculate the ratio of divergence angles. As the divergence angle of this type of alternate pattern changes in the sequence of *p*, *q*, −*p*, and −*q* (−180° < *p*, *q* ≤ 180°), |*q*|/|*p*| gives the ratio of the absolute values of divergence angles if |*p*| > |*q*|. Typical examples of phyllotactic patterns are marked with circled numbers in the color legend and their schematic diagrams are shown at the bottom.

### Numerical solution

To solve equations numerically, C++ with Visual C++ in Microsoft Visual Studio® 2019 was used for calculations.

## Results

### Morphological characterization of phyllotaxis of *C. megalobractea*

To characterize morphologically real costoid phyllotaxis, we performed anatomical analysis of the shoot apex of *C. megalobractea* using materials collected from adult vegetative plants (Fig 3). Similarly to the other members of Costaceae, *C. megalobractea* had obviously much smaller divergence angles, of which mean was about 54°, than those of common phyllotactic spirals (Fig 3A). The natural log of the plastochron ratio (*G*) (Richards, 1951) was about 0.62 (Fig 3C), which is very large comparable to the *G* values measured for distichous and spirodistichous patterns (Rutishauser, 1998).

**Fig 3.**
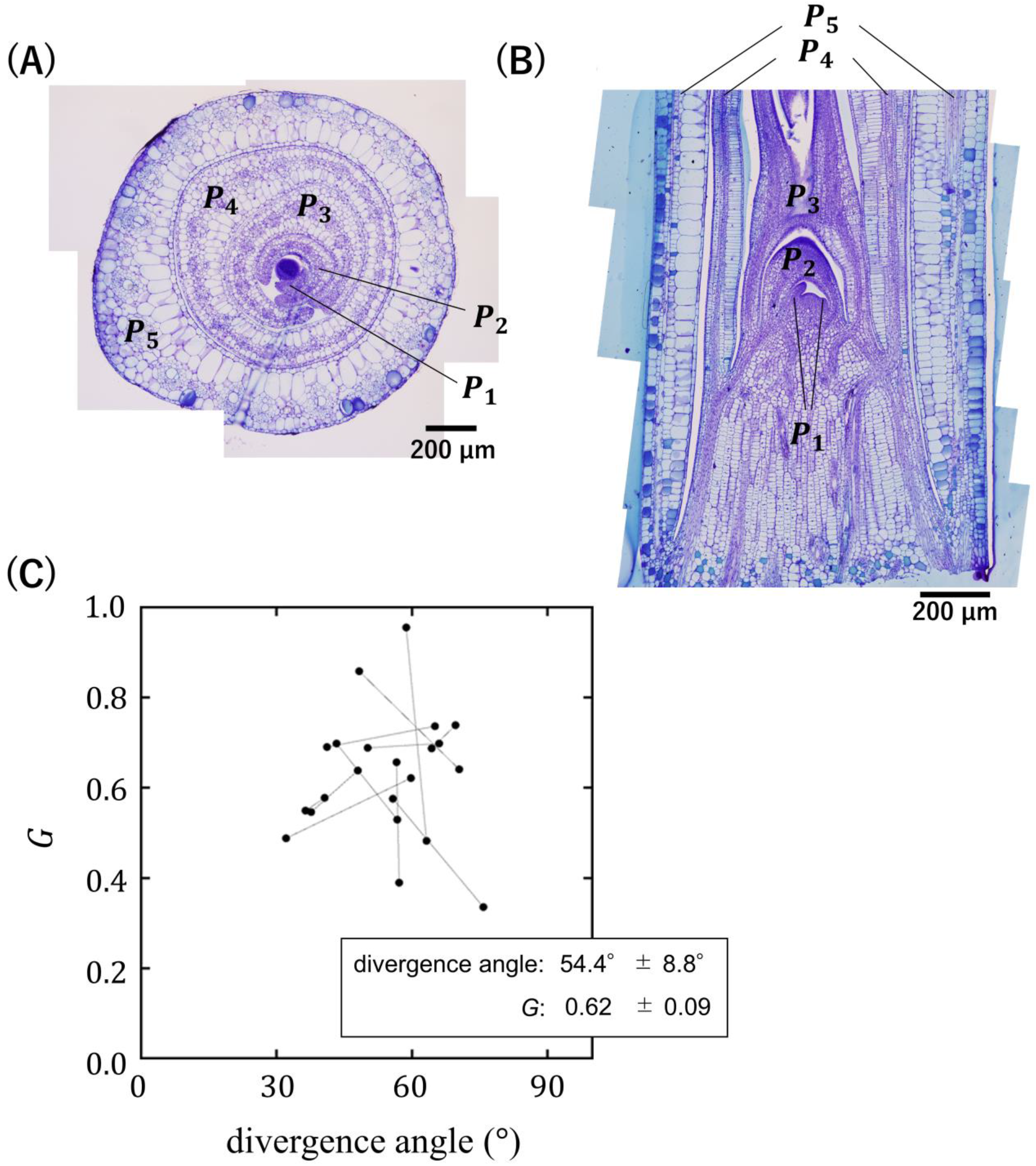
Costoid phyllotaxis in the vegetative shoot apex of *C. megalobractea*. (A) Transverse section. Leaf primordia are designated as *P*_1_, *P*_2_, *P*_3_, etc., with *P*_1_ being the youngest visible primordium. (B) Longitudinal section. (C) Divergence angles and *G* of *P*_1_ − *P*_2_ and *P*_2_ − *P*_3_ measured using the transverse sections. Points linked by a line represent data from the same sample.

Next, we inspected young seedlings of *C. megalobractea* to examine the early phyllotactic transition (Fig 4A). As previously reported in the other species of Costaceae (Weisse, 1932), the divergence angle of this plant decreased almost monotonically and converged quickly to a constant value (Fig 4B). The opposite positioning was not observed even between the cotyledon and the first true leaf (*L*_1_) or between *L*_1_ and the second true leaf (*L*_2_) (Fig 4A).

**Fig 4.**
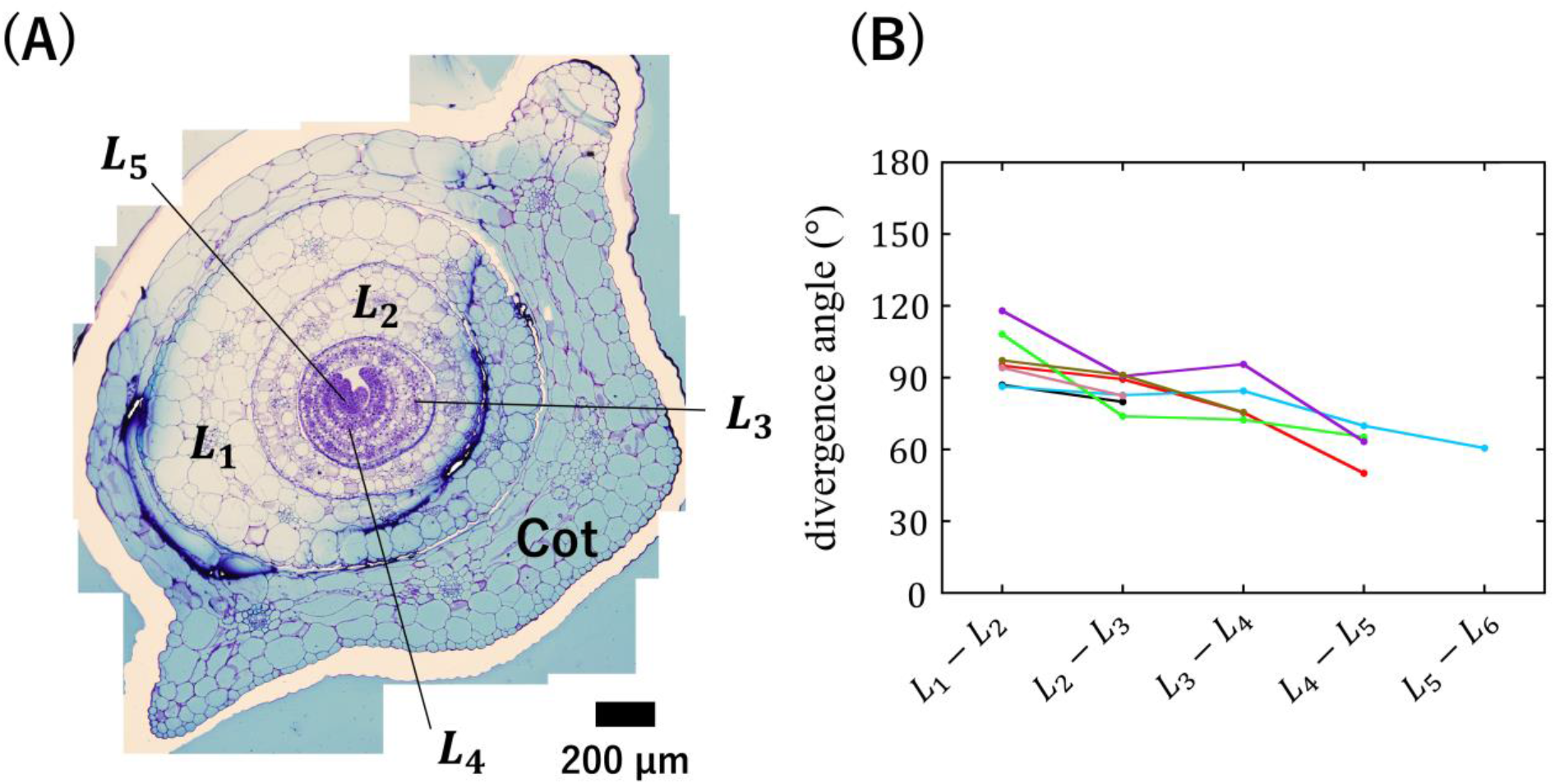
Early phyllotactic transition in the seedlings of *C. megalobractea*. (A) Transverse section of the shoot apex of a seedling. Leaf primordia are designated as *L*_1_, *L*_2_, *L*_3_, etc., with *L*_1_ being the oldest primordium. Cot indicates the cotyledon. (B) Divergence angles measured using the transversal sections of the seedling shoot apices with 7 samples indicated by different colors.

### Examination of the possibility of the occurrence of costoid phyllotaxis in the scheme of EDC models

As a preliminary step of mathematical model studies of costoid phyllotaxis, we examined EDC models for their ability of producing costoid phyllotaxis. We first searched for the conditions that generate steep spirals with relatively large plastochron ratios characteristic of the costoid pattern in the extensive computer simulations with EDC1, which allows only alternate primordium formation at a constant time interval (Yonekura et al., 2019). In the parameter space of EDC1, we identified a broad region generating such steep spirals, where the inhibitory power increases late and quickly (Fig S1A). Unlike real costoid phyllotaxis, however, the steep spiral generation by EDC1 did not show a monotonic decrease or quick convergence of the divergence angle: Instead, there was a long oscillatory phase including 0° sequences in the divergence angle before reaching a stable pattern (Fig S1A). Additionally, the parameter conditions for steep spiral generation were found to be close to the conditions that cause very strange alternate patterns with a multicycle change in the divergence angle (Fig S1B – D), which implies that these and neighboring conditions are quite unnatural. For all these reasons, we judged that EDC1 is inappropriate for the realistic production of costoid phyllotaxis.

We next examined whether costoid phyllotaxis can occur in the computer simulations with EDC2. An overall trend was identified in the alternate patterns generated by EDC2 that the plastochron ratio gets smaller as the divergence angle gets smaller. we confirmed that, according to this trend, at large plastochron ratios equivalent to that of real costoid phyllotaxis of *C. megalobractea*, only distichous phyllotaxis can be formed (Fig S2), which clearly excluded the occurrence of costoid phyllotaxis within the framework of EDC2.

Inspired by the idea of Hirmer (1922) that costoid phyllotaxis might be a kind of (spiro-)distichous phyllotaxis with one ortho/parastichy disappearing, we also considered the possibility that not all incipient primordia develop into visible primordia for the phyllotactic patterns generated by EDC2. When only every third incipient primordia become visible in Fibonacci spiral, the apparent divergence angle should be about 137.5° × 3 − 360° = 52.5°. When incipient primordia of spirodistichous phyllotaxis only alternately become visible, the apparent divergence angle should be about 360° − (145°~170°) × 2 = 70°~20°. In either case, the platochron ratio is sufficiently large and lies within the acceptable range. Thus, with respect to the divergence angle and the plastochron ratio, these patterns are similar to costoid phyllotaxis. We noticed, however, that they are distinct from real costoid phyllotaxis observed with *C. megalobractea* in the early phyllotactic transition: while the divergence angle monotonically decreased in real costoid phyllotaxis, the apparent divergence angles of the above patterns showed damped oscillation, reflecting the damped oscillation of the original spirals in the computer simulation with EDC2 (Figs S3 and S4). We therefore concluded that EDC2 cannot explain the generation of costoid phyllotaxis.

### Construction of a new model of phyllotactic pattern formation

The small divergence angle of costoid phyllotaxis means that the radial position of each leaf primordium is close to that of its preceding primordium, and the large plastochron ratio suggests that the positioning of the incipient primordium on the SAM periphery is seemingly influenced by only its immediately preceding primordium among the existing primordia because older primordia are already too far away from the SAM periphery. Taking these features as they are, we hypothesized that there may be some inductive effect, in addition to the inhibitory effect, from the preceding leaf primordium on new primordium formation. Based on this hypothesis, we constructed a new mathematical model that assumes both inductive and inhibitory fields. In the new model, an inductive filed created by the inductive power of each existing primordium is imposed on EDC2, and it is assumed that new primordium initiation is allowed only when the inductive and inhibitory field strengths are sufficiently high and sufficiently low, respectively.

The essential points of the new model are as follows.

1. The shoot apex is considered as a cone with an apical angle of *ψ*.
2. Each leaf primordium *L* emits an inductive power and an inhibitory power, which generates an inductive field and inhibitory field around it, respectively.
3. The inductive field and the inhibitory field strength declines as a function of the distance *d*.
4. The inductive power and the inhibitory power increase dependently on the primordial age *t*.
5. The formation of new primordia is restricted to the meristem periphery represented by the circle *M* with a distance of *R*_0_ from the conical vertex.
6. When the inductive field strength *Y* rises above a given threshold *Y_th_*, and the inhibitory field strength *S* falls below a given threshold *S_th_* somewhere on *M*, a new primordium is formed immediately at that point.
7. Primordia move away from the center of the shoot apex with a radial velocity that is proportional to the radial distance *r* due to the exponential growth of the shoot apex.

Positions on the conical surface are expressed in spherical coordinates 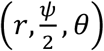.

Because of assumption 7, the distance from the center of the shoot apex to the *m*^th^ primordium on the conical surface (*r_m_*) is expressed with the time after its emergence *T_m_* and the initial radial velocity *V*_0_ as:

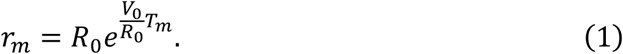

By using *t_m_* ≡ *T_m_ V*_0_/*R*_0_, a standardized age of the *m*^th^ primordium defined as the product of *T_m_* and the relative SAM growth rate *V*_0_/*R*_0_, *r_m_* is more simply expressed as:

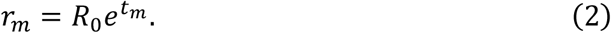

*t_m_* – *t*_*m*+1_ gives a standardized plastochron and it can be calculated from Eq 2 as:

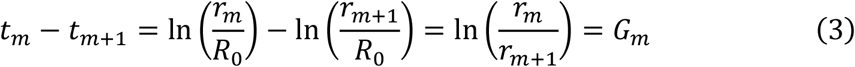

Thus, the standardized plastochron is equal to the natural log of the plastochron ratio and represented by *G_m_*.

The inductive field strength *Y*(*θ*) and inhibitory field strength *S*(*θ*) at the position 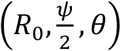 on *M* are calculated by summating inductive and inhibitory effects from all preceding primordia, respectively, as follows.

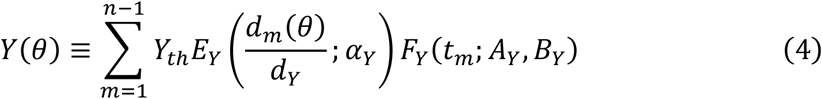

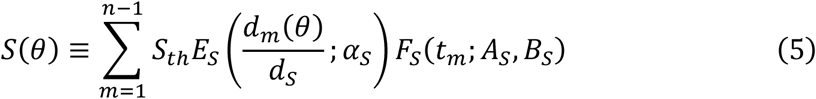

Here *d_m_* is a distance between the *m*-th primordium and the position 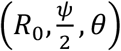, *d_Y_* and *d_S_* are the maximum distances within which an existing primordium induces a new primordium and which an existing primordium excludes a new primordium, respectively. *E* is a distance-dependent factor, which is defined as:

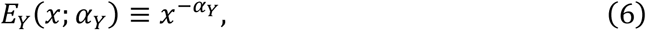

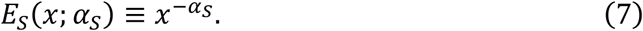

*F* is a primordial age-dependent factor, which is defined as:

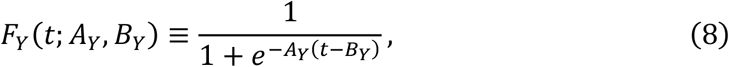

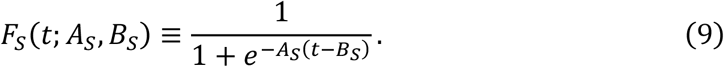

If *Y*(*θ*) > *Y_th_* and *S*(*θ*) < *S_th_*, a new primordium is placed at the position 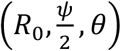. In this study, *Y_th_*, and *S_th_* are fixed to 1. *R*_0_ is also fixed to 1 unless otherwise indicated.

The new model is characterized by nine independent parameters; 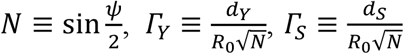, *α_Y_*, *α_S_*, *A_Y_*, *B_Y_*, *A_S_*, *B_S_*. These parameters represent the flatness of the shoot apex, the ratio of the maximal induction range to the SAM size, the ratio of the maximal inhibition range to the SAM size, the steepness of the decline of the inductive effect around the threshold, the steepness of the decline of the inhibitory effect around the threshold, the rate of the age-dependent increase of the inductive power, the timing of the increase of the inductive power, the rate of the age-dependent increase of the inhibitory power, and the timing of the increase of the inhibitory power, respectively.

### Generation of costoid phyllotaxis in computer simulation with the new model

Extensive computer simulations with the new model were conducted at *N* = 1/3 under a wide range of combinations of the other eight parameters (Figs S5 – S10). As a result, we detected the occurrence of steep spirals (green to cyan-green in these figures) in a particular range of conditions close to the conditions where no primordia were formed.

The relationships between the parameter conditions and the resultant patterns can be summarized as follows. Under such conditions that the increase of the inductive power is sufficiently fast and the inductive field strength is always larger than the threshold *Y_th_*, when and where the inhibitory field strength falls below the threshold *S_th_*, the patterns generated in the simulations with the new model are the same as generated by the EDC2 simulation (exemplified by Fig 6A), and thus costoid-phyllotaxis-like patterns are never produced. Under such conditions that the increase of the inductive power is very late and the inductive field strength is always smaller than *Y_th_* when and where the inhibitory field strength falls below *S_th_*, no new primordium formation takes place. Under intermediate conditions in which the increase of the inductive power is moderately late, steep spirals are generated in some cases (Fig 6B). These steep spirals were found to have not only small divergence angles but also large plastochron ratios (e.g., the divergence angle is 55.9° and the natural log of the plastochron ratio is 0.682 for the spiral shown in Fig 6B) and hence judged as representing costoid phyllotaxis. In simulations with varying the parameters related to the inductive field and with all other parameters fixed, there was an inverse correlation between the plastochron ratio and the divergence angle of the generated patterns (Fig S13).

**Fig 5.**
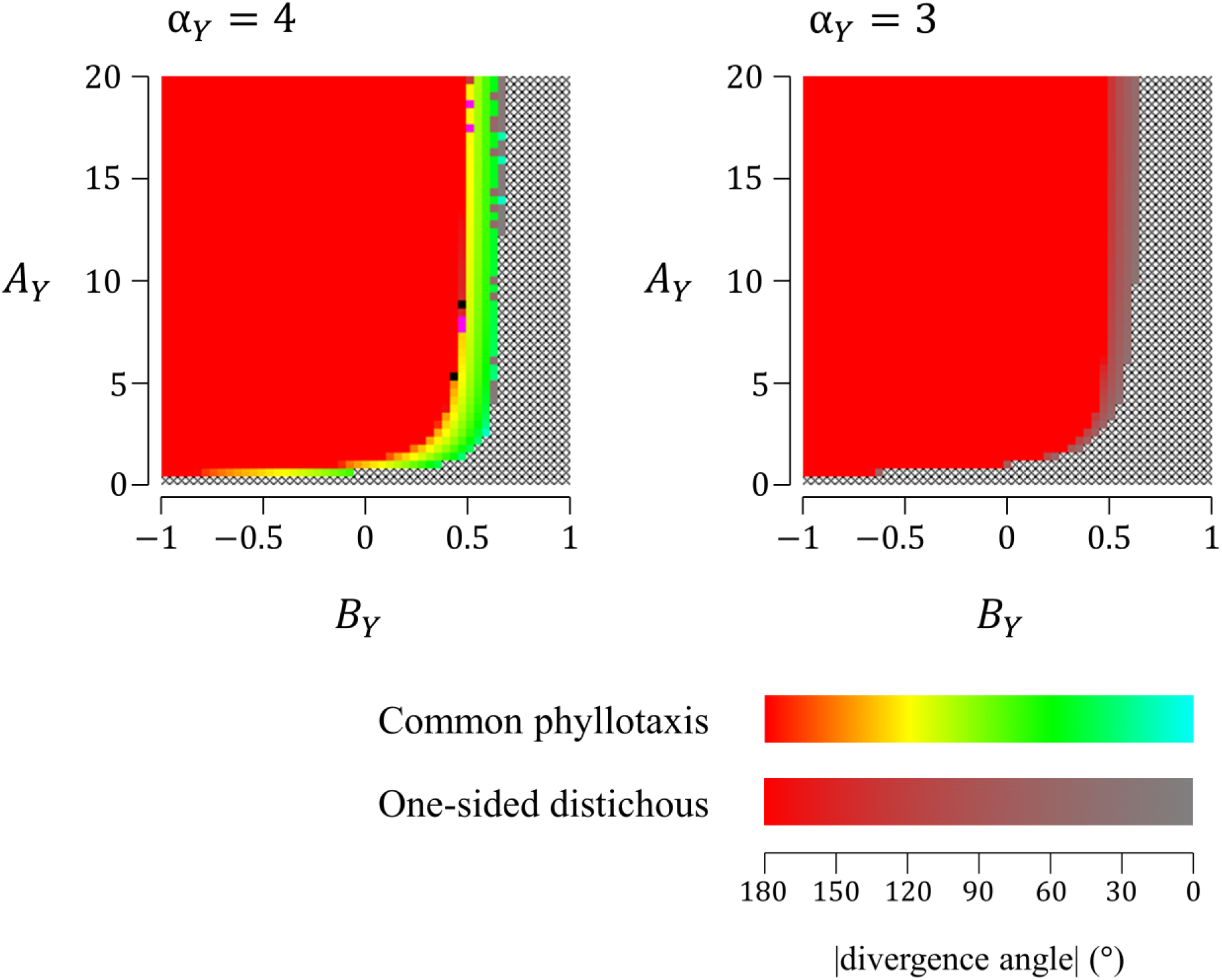
Distributions of phyllotactic patterns generated in the parameter space of the new model as influenced by *α_Y_*. Computer simulations using the new model were performed under various parameter settings (51 settings for 0 ≤ *A_Y_* ≤ 20, 51 settings for −1 ≤ *B_Y_* ≤ 1) with fixed parameters *N* = 1/3, *α_Y_* = 3 or 4, *α_S_* = 3, *Γ_Y_* = 3.5, *Γ_S_* = 3, *A_S_* = 10, and *B_S_* = 0, and the patterns obtained are displayed according to the color legend shown in Fig 2. Simulations were started by placing a single primordium on the SAM periphery.

**Fig 6.**
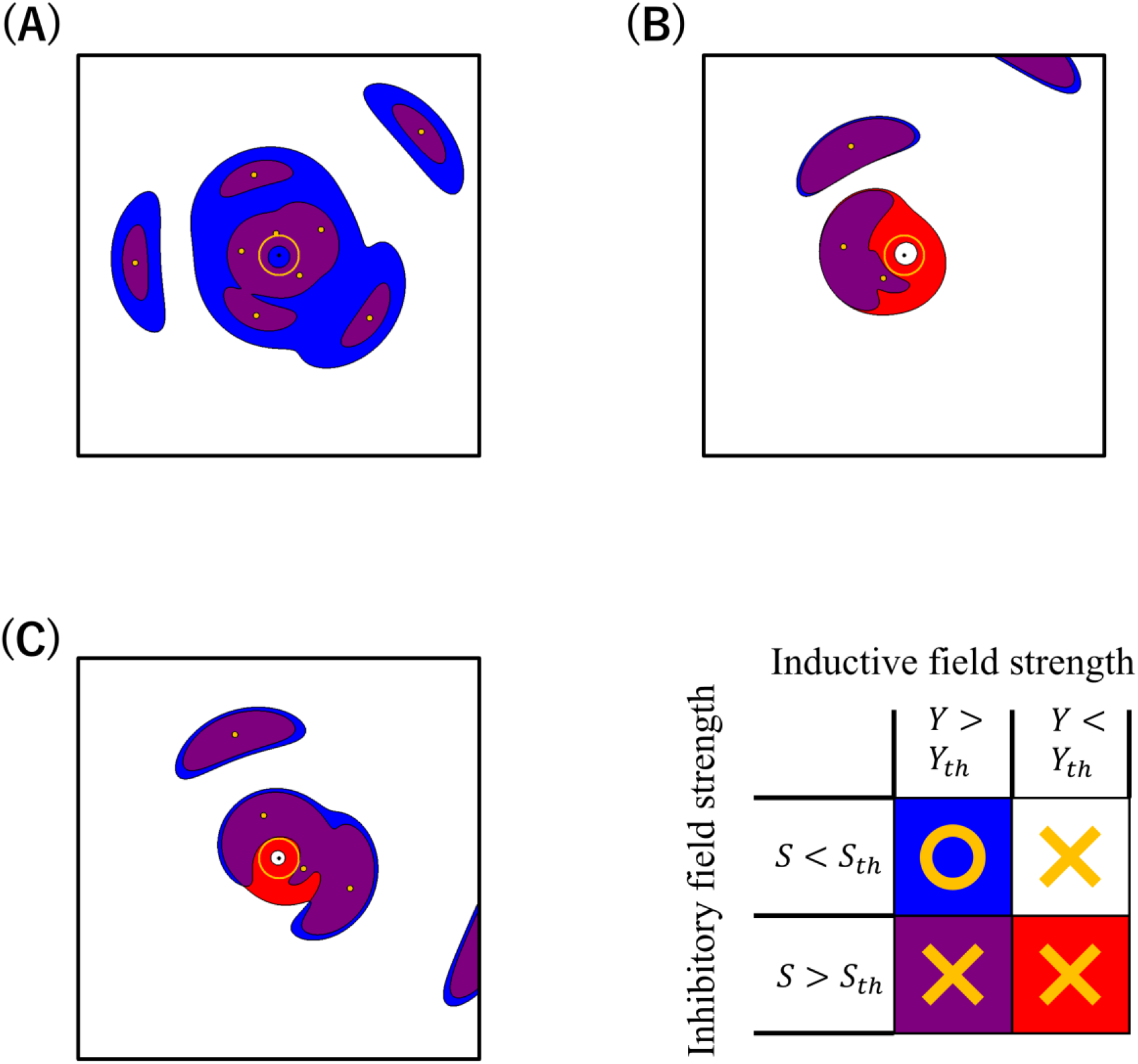
Phyllotactic patterns generated in computer simulations using the new model. Contour map of the inductive field strength *Y* and the inhibitory field strength *S* within the shoot apical region. Yellow circle indicates the periphery of the shoot apex, on which new primordia initiate. Yellow dots indicate the center of primordia. (A) Fibonacci spiral. Parameters were set to *N* = 1/3, *α_Y_* = 3, *α_S_* = 3, *Γ_Y_* = 3.5, *Γ_S_* = 1.9, *A_Y_* = 20, *B_Y_* = 0, *A_S_* = 10, and *B_S_* = 0. (B) Costoid phyllotaxis. Parameters were set to *N* = 1/3, *α_Y_* = 4, *α_S_* = 2, *Γ_Y_* = 3.5, *Γ_S_* = 3, *A_Y_* = 20, *B_Y_* = 0.64, *A_S_* = 10, and *B_S_* = 0. (C) One-sided distichous. Parameters were set to *N* = 1/3, *α_Y_* = 3, *α_S_* = 3, *Γ_Y_* = 3.5, *Γ_S_* = 3, *A_Y_* = 20, *B_Y_* = 0.52, *A_S_* = 10, and *B_S_* = 0.

We also found that it is necessary for the generation of the costoid pattern that the steepnesses of the distance-dependent decline of the inductive inhibitory effects satisfy *α_Y_* ≥ *α_S_*: otherwise, the delay of the increase of the inhibitory power resulted in the generation of the one-sided distichous pattern instead of the costoid patten (right panel of Fig 5, Fig 6C). One-sided distichy is a rare type of phyllotaxis, in which two orthostichies do not lie on a straight line but are angled to each other.

### Theoretical requirements for the generation of the costoid or one-sided distichous pattern in the new model

The results of the computer simulations with the new model indicated that the moderately late increase of the inductive power relative to the inhibitory power is important for the generation of the costoid or one-sided distichous pattern. From this finding, we presumed that it gives the requirements for the costoid or one-sided distichous pattern formation that the inductive field strength rises above the threshold at some point on the SAM periphery where the inhibitory field strength is below the threshold. To identify these requirements theoretically, we considered a simplified situation of the new model in which only one primordium of the standardized age *t*\* exists at 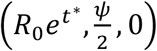 and both the inductive field strength and the inhibitory field strength reach their thresholds at the point 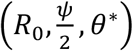 on the circle *M* (Fig S11). That is, the boundary of the induction range (within which the inductive field strength is larger than the threshold *Y_th_*) surrounding the single existing primordium meets the boundary of the inhibition range (within which the inhibitory field strength is larger than the threshold *S_th_*) surrounding the same primordium on *M*. As a new primordium arises at the position 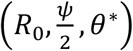 at the very moment in this situation, *t*\* and *θ*\* represent the standardized plastochron and the divergence angle, respectively. In this situation, the requirements for the existence of the solutions of *t*\* and *θ*\* are presumed to be equivalent to the requirements for the generation of the costoid or one-sided distichous patterns.

The above situation can be described as follows by comparing Eq 4 to Eq 5:

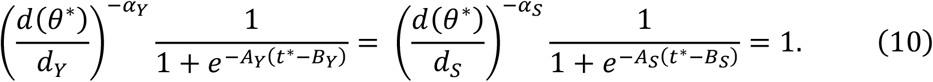

Using *Γ_Y_* and *Γ_S_* to eliminate *d_Y_* and *d_S_*, Eq 10 can be described as follows:

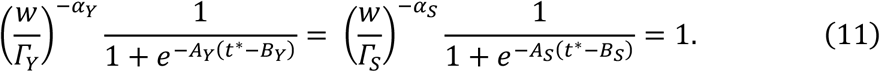

Here *w* is defined as:

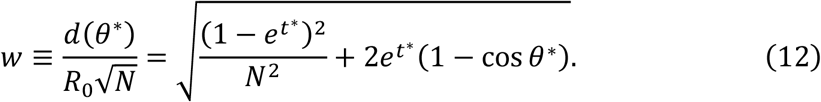

*t*\* is obtained as the solution of the following equation derived from Eq 11:

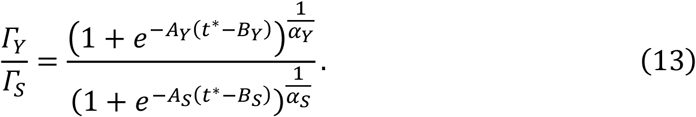

After we solved Eq 13 for *t*\* numerically, we calculated *w*:

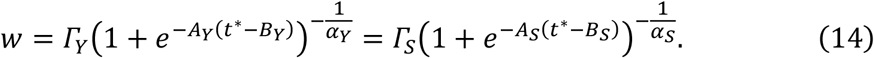

Then we calculated *θ*\* using the following equation derived from Eq 12:

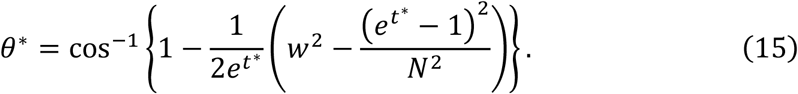

The solutions of *t*\* and *θ*\* were plotted, if they exist, against *B_Y_*, and were compared with *G* and the divergence angle of the patterns generated by the new model simulations with varying *B_Y_* (Fig S12). This comparison indicated that the range of *B_Y_* where the solutions of *t*\* and *θ*\* exist is almost correspondent to the *B_y_* range where the new model produced the costoid or one-sided distichous pattern and that, in this range, *t*\* and *θ*\* fit well to the *G* value and the divergence angle of the model-generated patterns, respectively. The good agreement supported our presumption regarding the requirements for the generation of costoid or one-sided distichous pattern. Now the following explanation can be deduced. When *B_Y_* is enough small, the induction range always encompasses the inhibition range, and in such case, common types of phyllotaxis are generated. When *B_Y_* is moderately larger, the induction range is encompassed by the inhibition range at first and later it expands beyond the inhibition range somewhere on *M*, and in such case, costoid phyllotaxis or one-sided distichous patterns are generated. When *B_Y_* is too large, the induction range is always encompassed by the inhibition range, and such case can never produce new primordia.

More accurately, however, there are some differences between the solutions obtained from the simplified situation and the simulation results. The *t*\* values are slightly smaller than the *G* values (upper panels of Fig S12), which should reflect the small effect from the second youngest primordium and/or older primordia. Moreover, the *B_Y_* range generating costoid or one-sided distichous patterns in the simulation is a little different from the *B_Y_* range for the existence of the solutions of *t*\* and *θ*\* (Fig S12). Such discrepancy is seen in the range of *B_Y_* where *t*\* is smaller than *G_S_*, the *G* value calculated from the simulation without considering the inductive effect (yellow zone in Fig S12), which is also attributable to the effect of the second youngest primordium and/or older primordium.

From the above consideration on the simplified situation of the new model, we found a straightforward relationship between the SAM size *R*_0_ and the divergence angle *θ*\*. By substituting the definitions 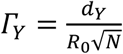 and 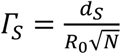 into Eq 12 and Eq 14, we have:

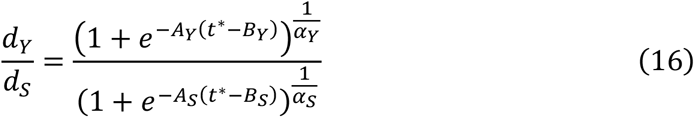

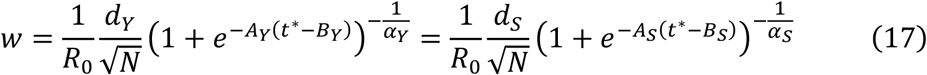

Eq 16 indicates that *t*\* is independent from *R*_0_ and is constant when the other parameters are fixed. *w* can be written as:

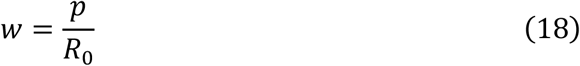

where 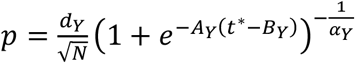 is also independent from *R*_0_ and is constant.

Using Eq 18, Eq 15 can be modified to:

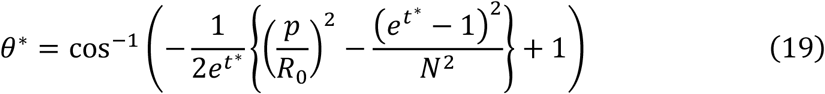

Eq 19 shows the obvious relationship that *θ*\* get smaller when *R*_0_ get larger and when all other parameters are fixed to constant values (Fig S13).

### Early changes in the divergence angle of costoid phyllotaxis in the new model

From the negative correlation between the SAM size and the divergence angle in Eq 19, the possibility arose that the monotonical decrease of the divergence angle in early leaves of the seedlings of *C. megalobractea* is associated with the increase of the SAM size, since the enlargement of SAM during seedling development has been often observed in many plants (e.g. *Arabidopsis thaliana* in Medford et al., 1992; *Zea mays* in Thompson et al., 2014).

Douady and Couder (1996c) already dealt with the enlargement of SAM, by describing the effect of the time-dependent change of the SAM size on *Γ_S_* the as:

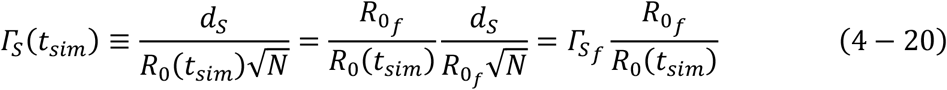

where *R*_0_*f*__ indicates the final SAM size at *t* → ∞, and 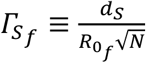. Thus, the SAM enlargement was introduced as the function of *Γ_S_*(*t_sim_*) (Douady and Couder, 1996c):

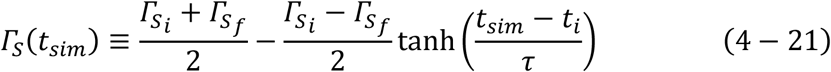

where *t_sim_* is the passing time in the simulation *t_i_* and *τ* represent the timing and gentleness of the SAM enlargement, respectively. *Γ*_*S*_*i*__ indicates the limiting value of the *Γ_S_*(*t_sim_*) at *t* → – ∞.

We applied this way of dealing with the SAM enlargement to the new model, with introducing an additional parameter *Γ*_*Y*_*f*__ to describe the effect from the time-dependent change of the SAM size on the inductive field. By substituting Eq 20, *Γ_y_*(*t_sim_*) is written as:

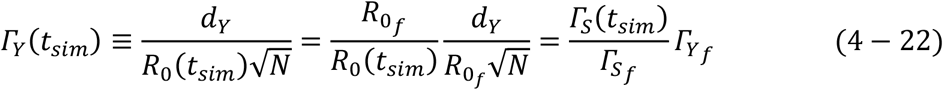

Computer simulation with the new model assuming a small increase in the SAM size showed a monotonical decrease of the divergence angle in the costoid pattern (Fig 7). This transition fitted well to the observations made with seedlings of *C. megalobractea*, which reinforced that the new model is plausible as a model of comprehensive generation of phyllotactic patterns including realistic costoid phyllotaxis.

**Fig 7.**
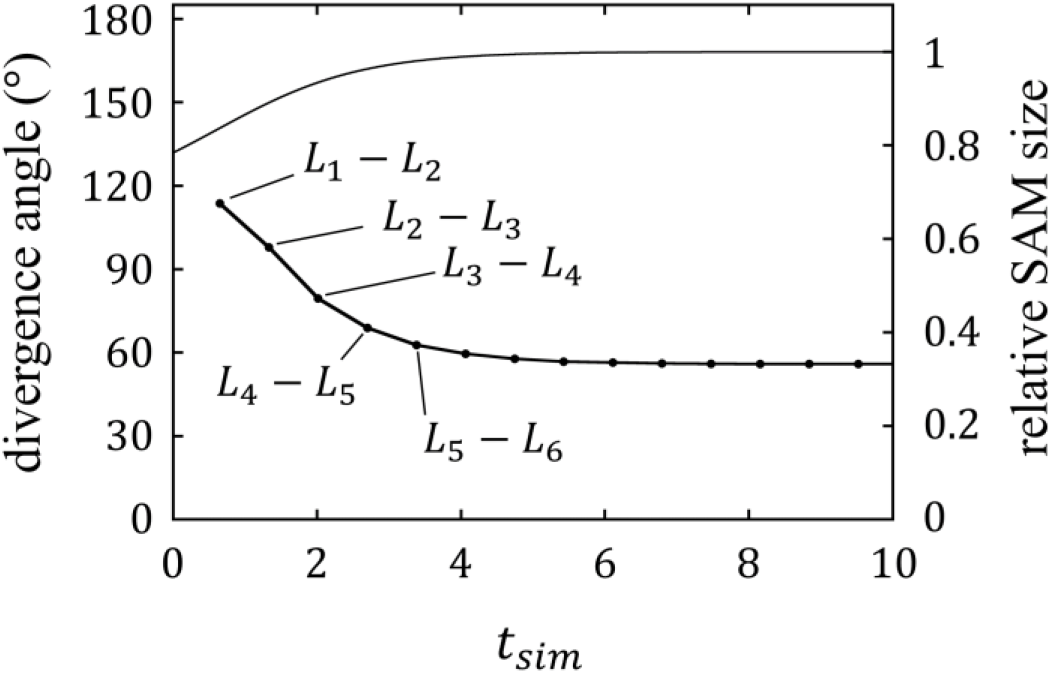
Early phyllotactic transition of costoid phyllotaxis obtained from the computer simulation using the new model with the SAM enlargement. The line chart shows the divergence angles between *L_n_* and *L*_*n*+1_ against the timing of the emergence of *L*_*n*+1_. The first true leaf primordium *L*_1_ was set to emerge at *t_sim_* = 0. The curve indicates the ratio of the SAM size to the final SAM size *R*_0_*f*__. Parameters were set to *N* = 1/3, *α_Y_* = 4, *α_S_* = 2, *Γ_Y_* = 3.5, *Γ*_*S*_*i*__ = 4.5, *Γ*_*S*_*f*__ = 3, *A_Y_* = 20, *B_Y_* = 0.64, *A_S_* = 10, and *B_S_* = 0.

## Discussion

Costoid phyllotaxis showing spiromonostichy unique to Costaceae is famous as the “genuine puzzle” in the developmental studies of phyllotaxis because its peculiar morphological characters, such as a steep spiral with a small divergence angle and a large plastchron ratio, challenge Hofmeister’s axiom that has served as the basis of all mechanistic models of phyllotactic pattern formation (Kirchoff and Rutishauser, 1990). Here we attempted to solve this puzzle by mathematical model analysis with EDC2 as a starting point. We first compared the steep spiral patterns generated in the computer simulations with EDC2 to real costoid phyllotaxis that we observed in the shoot apex of *C. megalobractea*. The mean divergence angle of costoid phyllotaxis of this plant in the adult vegetative phase was about 54°. Comparable small divergence angles are found in only few plants outside Costaceae (Rutishauser, 1998; Zagórska-Marek et al., 2021). *Picea abies* is one of such exceptional plants and was reported to have a divergence angle of 56.4° (Rutishauser, 1998). The plastochron ratio in *C. megalobractea* was, however, much larger than that in *P. abies*. The natural log of the Richards’s plastochron ratio (*G*) of *C. megalobractea* was about 0.62, while *G* of *P. abies* is about 0.012. The large plastochron ratio measured in *C. megalobractea* is almost the same as the plastochron ratios of distichous and spirodistichous phyllotaxes (Rutishauser, 1998). Thus, the large plastochron ratio as well as the small divergence angle characterize costoid phyllotaxis and distinguish it from the other types of phyllotaxis. In contrast, among the phyllotactic patterns generated by EDC2, such large plastochron ratios were restricted to the distichous patterns and were not compatible with steep spirals with small divergence angles. We also considered the possibility that not all incipient primordia only grow into visible primordia. This scheme can produce apparently steep spirals having a large plastochron ratio but was found to contradict the early phyllotactic transition in the seedlings of *C. megalobractea*. These analyses led to the conclusion that EDC2 is not sufficient for the generation of costoid phyllotaxis.

Based on the morphological features of costoid phyllotaxis, we then hypothesized some effect of the existing leaf primordia to induce primordium initiation in their vicinity. we imposed this hypothetical inductive effect on EDC2 to construct the new model. Computer simulations using the new model successfully generated the costoid pattern, which matches real costoid phyllotaxis in several aspects, in addition to all phyllotactic patterns generated in EDC2. When the age-dependent increase of the inductive power occurs early and rapidly compared to the age-dependent increase of the inhibitory power, the new model is substantially not different from EDC2. When the age-dependent increase of the inductive power is too slow, no primordia are initiated in the new model. In a particular range of parameter settings between these two conditions, where the age-dependent increase of the inductive power is moderately slow, costoid phyllotaxis is generated.

Notably, the new model can generate not only costoid phyllotaxis but also one-sided distichy in similar conditions. One-sided distichous phyllotaxis, characterized by its primordium initiation position on two angled orthostichies, is one of rare types of phyllotaxis that have never been explained by the previously proposed mechanisms. The one-sided distichous pattern was reported for the inflorescence of *Thalia* and *Ctenanthe* (Marantaceae) and the rhizome of *Chamaecostus cuspidatus* (Costaceae) (Kirchoff, 1986; Schumann, 1902; Kirchoff and Rutishauser, 1990; Schüepp, 1928). As all these plants belong to Zingiberales, both costoid phyllotaxis and one-sided distichous phyllotaxis are characteristic of this plant group, implying some relation between them. The adjacent occurrence of these uncommon phyllotaxes in the parameter space of the new model may account for their close relationship and restriction to Zingiberales and is regarded as very supportive of the validity of the new model.

Which costoid phyllotaxis or one-sided distichous phyllotaxis is generated in the new model simulation depended on which parameter *α_Y_* or *α_S_* is larger. This result can be interpreted from the influence of these parameters on the inductive and inhibitory effects from the second youngest primordium *P*_2_ to the SAM periphery: if *α_Y_* > *α_S_*, the inhibitory effect from *P*_2_ is stronger than its inductive effect and a new leaf primordium arises at the flank of the youngest primordium *P*_1_ on the side distal to *P*_2_, leading to the generation of costoid phyllotaxis, otherwise a new primordium arises on the side proximal to *P*_2_, leading to the generation of one-sided distichous phyllotaxis

Theoretical considerations of the simplified situation of the new model in which only one preceding primordium exists revealed the principle of the mechanism working for the generation of the costoid and one-sided distichous patterns. According to this analysis, the costoid or one-sided distichous patterning requires that the induction range is encompassed by the inhibition range at first and later expands beyond the inhibition range somewhere on the SAM periphery. In this case, new primordium initiation takes place when and where the boundary of the induction range meets the boundary of the inhibition range on the SAM periphery.

Theoretical analysis also disclosed a negative relationship between the divergence angle and the SAM size in the costoid and one-sided distichous patterns generated in the new model. Based on this relationship, the early phyllotactic transition in the costoid pattern was assessed by computer simulations using the new model with assuming SAM enlargement. The result showed a monotonic decrease of the divergence angle, as observed in the seedlings of *C. megalobractea*. Although there are no data available for the SAM growth of *Costus*, the SAM enlargement assumed in this simulation was at 25% level of the SAM growth measured in *Zea mays* (Thompson et al., 2014) and does not seem unusual. Thus, the new model can reasonably explain the early phyllotactic transition in the seedlings of *C. megalobractea*. Collectively, these studies indicate that the new model reflects the actual mechanism of leaf primordium positioning and that the inductive field hypothesized in the new model is truly involved in phyllotactic pattern formation and particularly important for costoid and one-sided distichous phyllotaxes in Zingiberales.

In spite of the drastic modification of the model with the introduction of the hypothetical inductive effect, there are limited changes in the distribution of phyllotactic patterns in the parameter space between EDC2 and the new models: substantially the only difference is the occurrence of the costoid and one-sided distichous patterns in the new model within a narrow range at the marginal conditions, outside which no primordium initiation is allowed. It may be the reason why plants having costoid phyllotaxis are restricted to Costaceae that, without luckily acquiring some strict regulation, plants in this narrow range of parameters would easily lose leaves by a small change in the parameters and be eliminated by natural selection.

Molecular biological studies have demonstrated that leaf primordium initiation is controlled by the plant hormone auxin and its polar transport (Reinhardt et al., 2000; 2003; Benková et al., 2003). The current view on this control is that the positive feedback loop between auxin gradient and auxin polar transport by the auxin efflux carrier PIN1 creates auxin convergence, which triggers leaf primordium initiation (Smith et al., 2006; Jönsson et al., 2006). In this scheme, the inhibitory effect of the existing primordia on the vicinal formation of a new primordium is correspondent to the auxin depletion by drainage into the existing auxin convergence from its vicinity (Mirabet et al., 2012). On the other hand, leaf primordia and young leaves are known to be major sources of auxin in the shoot apex (Cheng et al., 2007; Galvan-Ampudia, C.S. et al., 2020). Furthermore, it was shown that the auxin biosynthesis enzyme YUCCA is necessary for initiation of shoot lateral organs in Arabidopsis and that YUC1 and YUC4, major members of YUCCA, are expressed mainly in lateral organ primordia in the shoot apex (Cheng et al., 2007). Most simply thinking from these roles of auxin, the possibility can be considered that, as well as the inhibitory effect, the inductive effect hypothesized in the new model may be mediated by auxin. Analysis of the dynamics of auxin biosynthesis and auxin transport with plants Costaceae would be important as the first step of the molecular characterization of costoid phyllotactic patterning and the new model.

