## Supplementary Figures for "A new mathematical model of phyllotaxis to solve the genuine puzzle spiromonostichy"

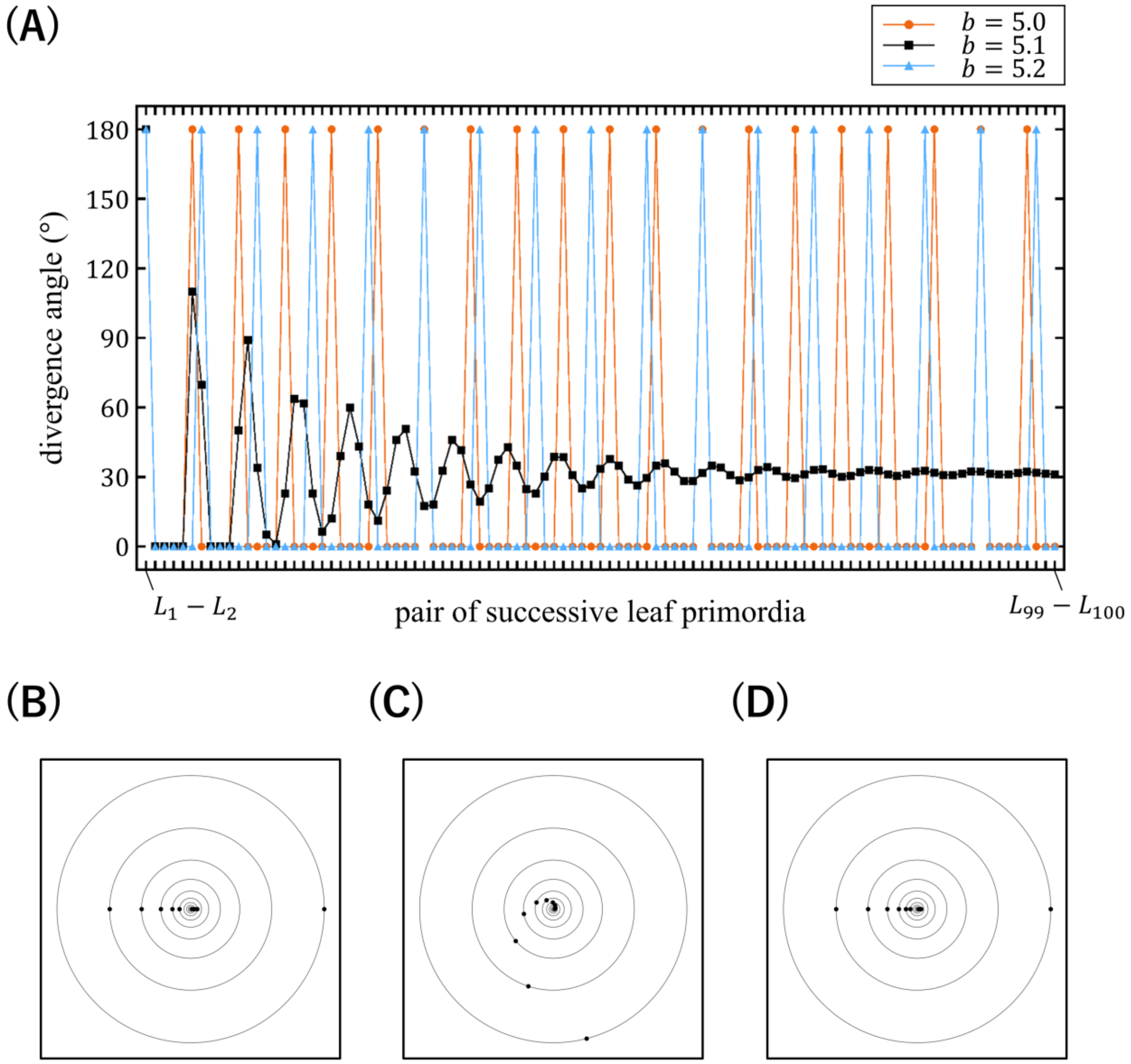


**Fig S1. Steep spiral pattern generated in the computer simulations with EDC1**

(A) Changes in the divergence angle in the patterns obtained by the computer simulation using EDC1 under slightly different settings of $b$ with fixed parameters $G = 0.5$, $a = 10$, and $\eta = 2$. (B) Five-cycle alternate pattern generated at $b = 0.50$. (C) Steep spiral pattern generated at $b = 0.51$. (D) Six-cycle alternate pattern generated at $b = 0.52$.

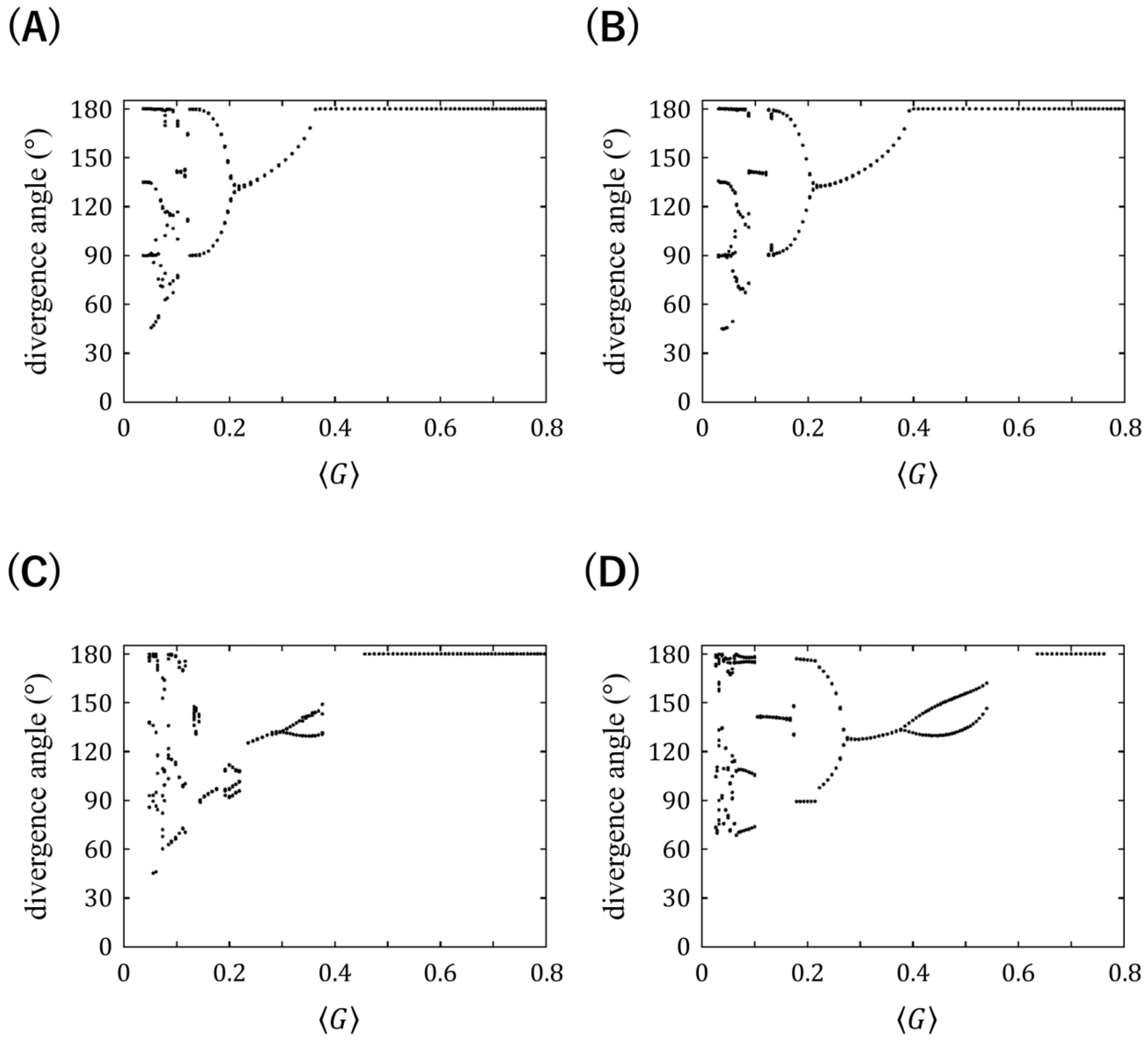


**Fig S2. Characteristics of phyllotactic patterns generated in the computer simulations with EDC2**

For each of the phyllotactic patterns generated in the computer simulations with EDC2, the divergence angle is plotted against the averaged natural log of the plastochron ratio (average of standardized plastochrons) $\langle G \rangle$. Data were obtained from the results shown in Fig 2-19, where simulations were performed under 101 settings of $\Gamma$ ($1 \leq \Gamma \leq 3$) at $N = 1/3$, $A = 5$ or $10$, $A \times B = 3$, and $\alpha = 1$ or $4$.

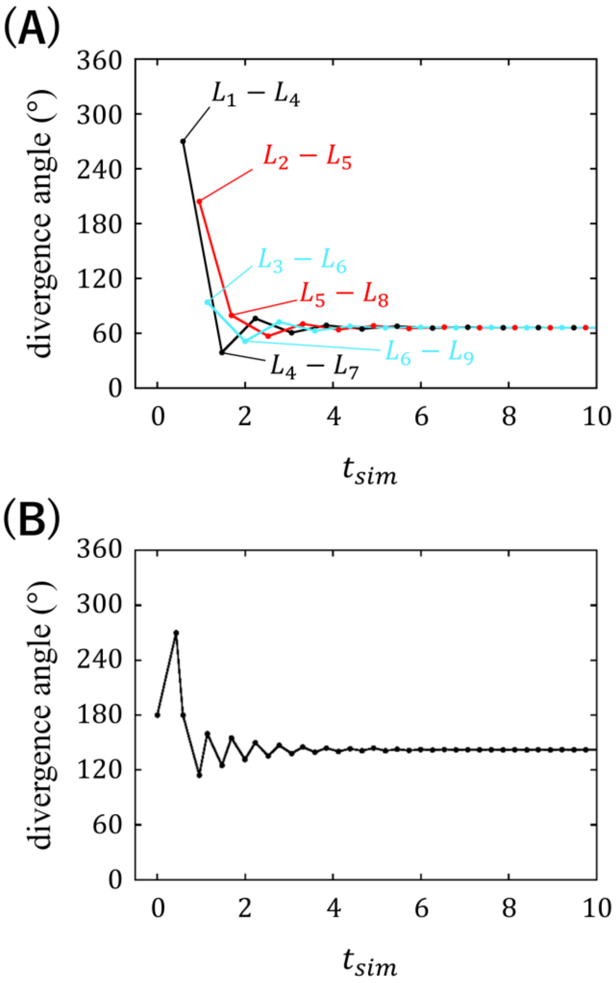


**Fig S3. Changes in the divergence angle of the Fibonacci spiral generated in the computer simulation with EDC2**

Fibonacci spiral was obtained by the computer simulation using EDC2 with parameters $N = 1/3$, $\Gamma = 1.9$, $\alpha = 3$, $A = 10$, and $B = 0$. (A) Changes in the divergence angle between the pair of primordia $L_{3n-2}$ and $L_{3n+1}$, $L_{3n-1}$ and $L_{3n+2}$, or $L_{3n}$ and $L_{3n+3}$, which represents the unusual situation where only every third incipient primordia develop into visible primordia. (B) Changes in the divergence angle between the primordia $L_n$ and $L_{n+1}$, which represents the normal situation where only all incipient primordia develop into visible primordia.

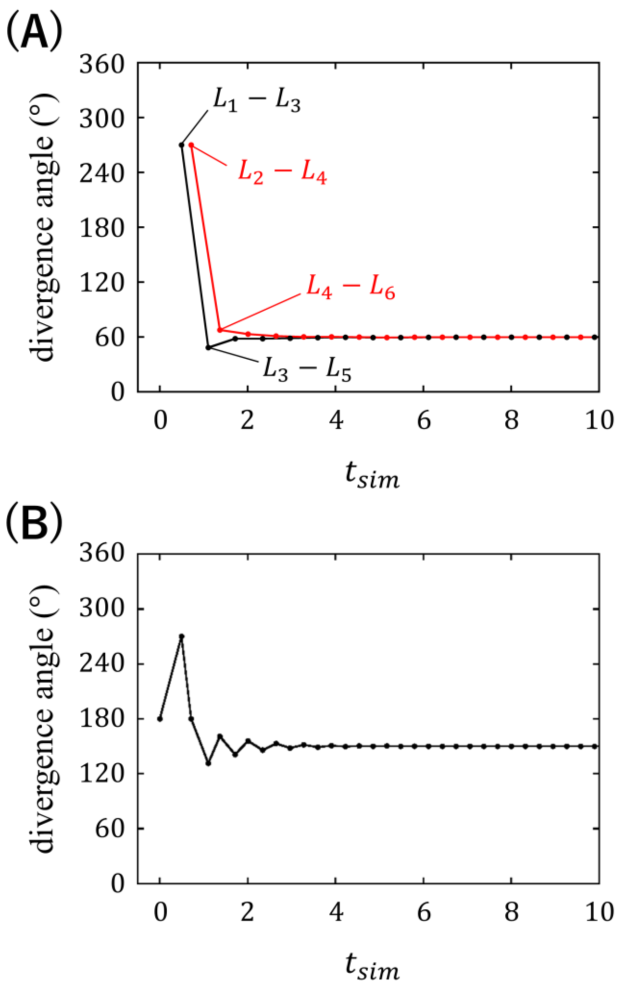


**Fig S4. Changes in the divergence angle of the spirodistichy generated in the computer simulations with EDC2**

Spirodistichy was obtained by the computer simulation using EDC2 with parameters $N = 1/3,\ \Gamma = 2.1,\ \alpha = 3,\ A = 10$, and $B = 0$. (A) Changes in the divergence angle between the pair of primordia $L_{2n-1}$ and $L_{2n+1}$ or $L_{2n}$ and $L_{2n+2}$, which represents the unusual situation where only every second incipient primordia develop into visible primordia. (B) Changes in the divergence angle between the primordia $L_n$ and $L_{n+1}$, which represents the normal situation where only all incipient primordia develop into visible primordia.

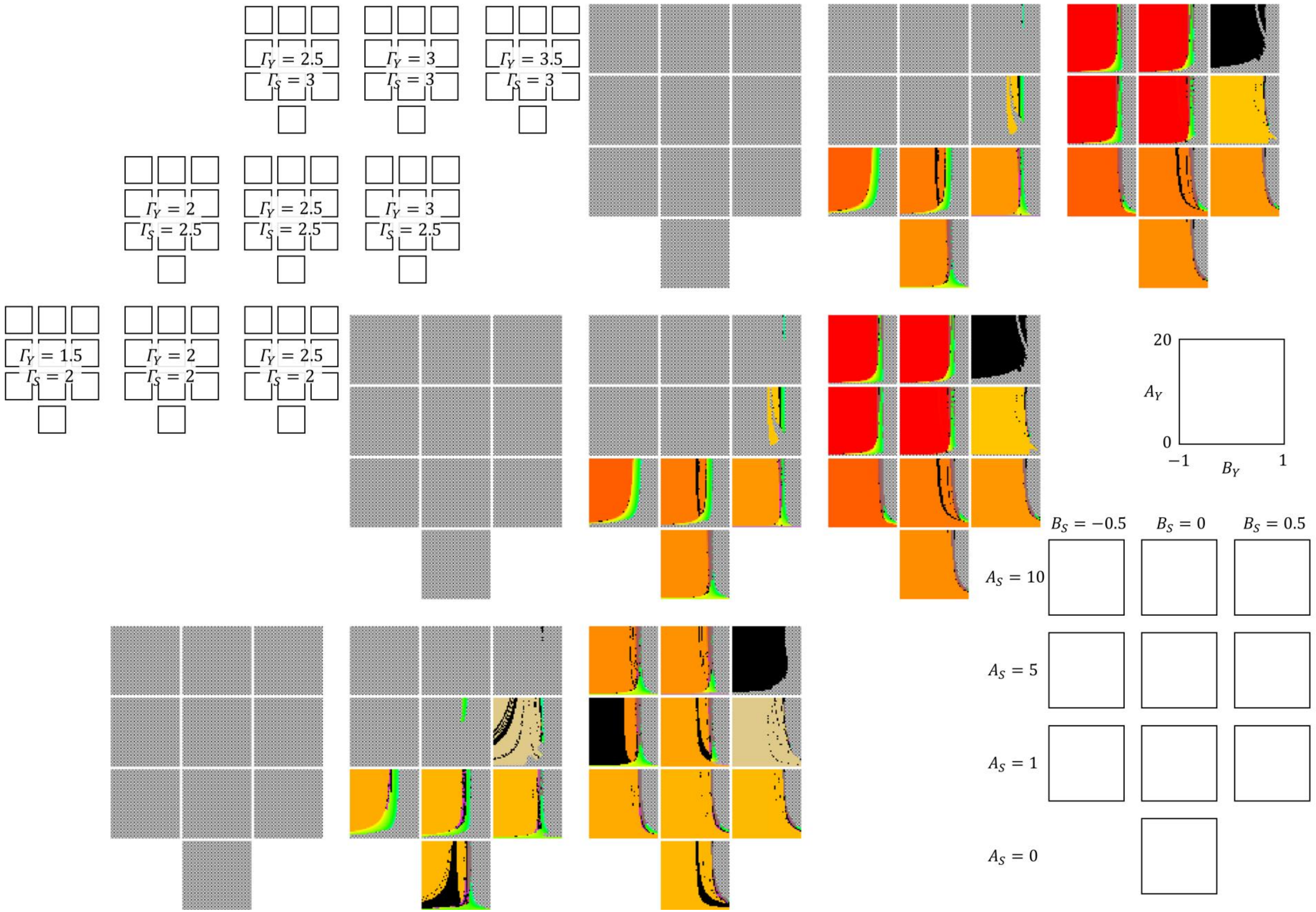

$\Gamma_Y = 2.5$
$\Gamma_S = 3$
$\Gamma_Y = 3$
$\Gamma_S = 3$
$\Gamma_Y = 3.5$
$\Gamma_S = 3$
$\Gamma_Y = 2$
$\Gamma_S = 2.5$
$\Gamma_Y = 2.5$
$\Gamma_S = 2.5$
$\Gamma_Y = 3$
$\Gamma_S = 2.5$
$\Gamma_Y = 1.5$
$\Gamma_S = 2$
$\Gamma_Y = 2$
$\Gamma_S = 2$
$\Gamma_Y = 2.5$
$\Gamma_S = 2$
20
$A_Y$
0
−1
$B_Y$
1
$B_S = -0.5$
$B_S = 0$
$B_S = 0.5$
$A_S = 10$
$A_S = 5$
$A_S = 1$
$A_S = 0$

**Fig S5. Computer simulations using the new model over a wide range of combinations of seven parameters focused on the inductive effect (1)**

Computer simulations were performed using the new model under various settings of eight parameters, $51$ settings for $A_Y$ $(0 \leq A_Y \leq 20)$, $51$ settings for $B_Y$ $(-1 \leq B_Y \leq 1)$, $9$ settings for $\Gamma_Y$ and $\Gamma_S$ $((\Gamma_Y, \Gamma_S) = (1.5, 2)$, $(2, 2)$, $(2.5, 2)$, $(2, 2.5)$, $(2.5, 2.5)$, $(3, 2.5)$, $(2.5, 3)$, $(3, 3)$, or $(3, 3.5))$, $4$ settings for $A_S$ ($A_S = 0$, $1$, $5$, or $10$), and $3$ settings for $B_S$ ($B_S = 0$, $-0.5$, or $0.5$). $N$, $\alpha_Y$, and $\alpha_S$ were fixed to $1/3$, $4$, and $3$, respectively. The patterns obtained are displayed in the $A_Y B_Y$ space according to the color legend shown in Fig 2. Simulations were started by placing a single primordium.

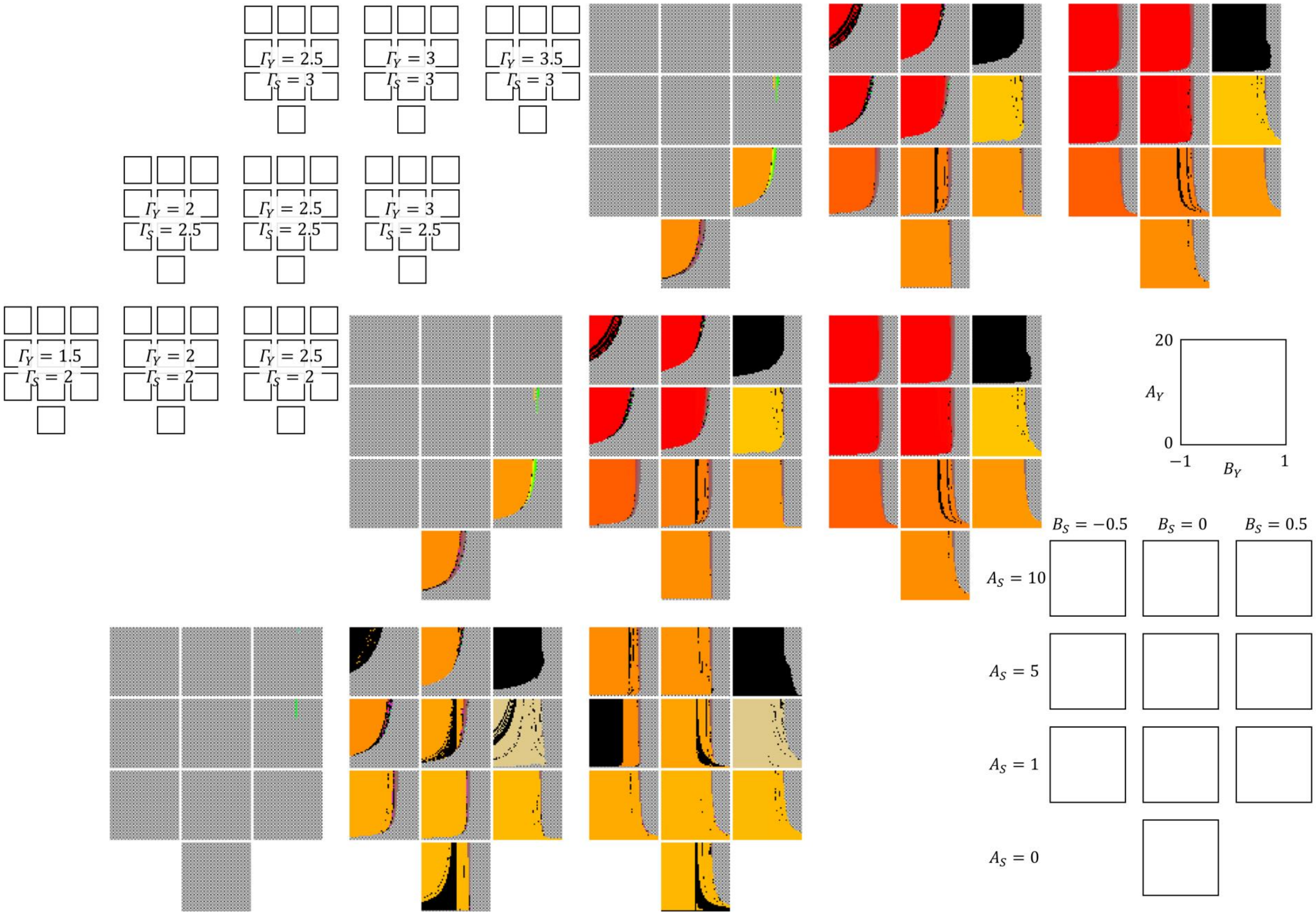

$\Gamma_Y = 2.5$ $\Gamma_S = 3$
$\Gamma_Y = 3$ $\Gamma_S = 3$
$\Gamma_Y = 3.5$ $\Gamma_S = 3$
$\Gamma_Y = 2$ $\Gamma_S = 2.5$
$\Gamma_Y = 2.5$ $\Gamma_S = 2.5$
$\Gamma_Y = 3$ $\Gamma_S = 2.5$
$\Gamma_Y = 1.5$ $\Gamma_S = 2$
$\Gamma_Y = 2$ $\Gamma_S = 2$
$\Gamma_Y = 2.5$ $\Gamma_S = 2$
20
$A_Y$
0
−1
$B_Y$
1
$B_S = -0.5$
$B_S = 0$
$B_S = 0.5$
$A_S = 10$
$A_S = 5$
$A_S = 1$
$A_S = 0$

**Fig S6. Computer simulations using the new model over a wide range of combinations of seven parameters focused on the inductive effect (2)**

Computer simulations were performed using the new model under various settings of eight parameters, $51$ settings for $A_Y$ $(0 \leq A_Y \leq 20)$, $51$ settings for $B_Y$ $(-1 \leq B_Y \leq 1)$, $9$ settings for $\Gamma_Y$ and $\Gamma_S$ $((\Gamma_Y, \Gamma_S) = (1.5, 2)$, $(2, 2)$, $(2.5, 2)$, $(2, 2.5)$, $(2.5, 2.5)$, $(3, 2.5)$, $(2.5, 3)$, $(3, 3)$, or $(3, 3.5))$, $4$ settings for $A_S$ ($A_S = 0$, $1$, $5$, or $10$), and $3$ settings for $B_S$ ($B_S = 0$, $-0.5$, or $0.5$). $N$, $\alpha_Y$, and $\alpha_S$ were fixed to $1/3$, $3$, and $3$, respectively. The patterns obtained are displayed in the $A_Y B_Y$ space according to the color legend shown in Fig 2. Simulations were started by placing a single primordium.

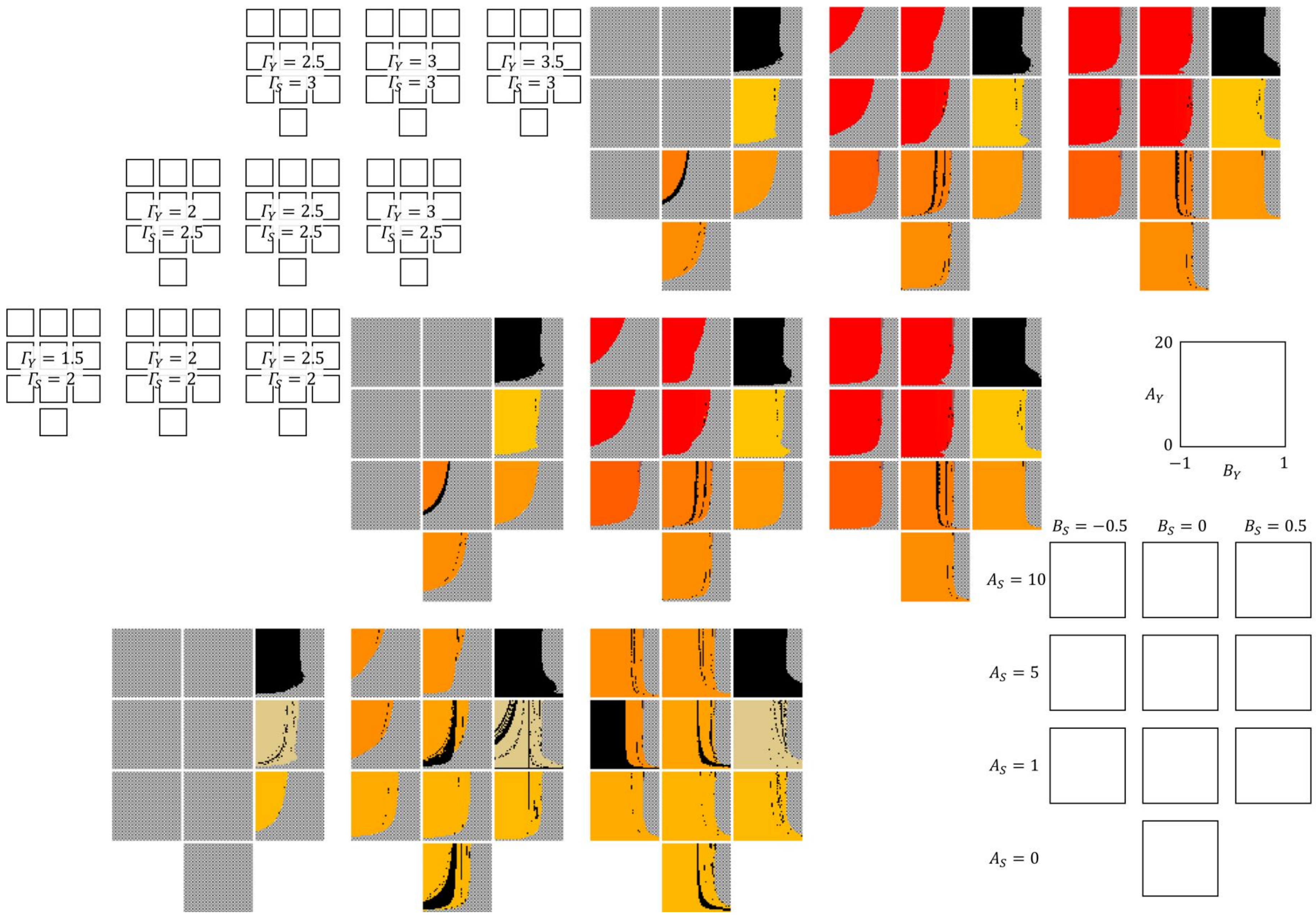
$\Gamma_Y = 2.5$
$\Gamma_S = 3$
$\Gamma_Y = 3$
$\Gamma_S = 3$
$\Gamma_Y = 3.5$
$\Gamma_S = 3$
$\Gamma_Y = 2$
$\Gamma_S = 2.5$
$\Gamma_Y = 2.5$
$\Gamma_S = 2.5$
$\Gamma_Y = 3$
$\Gamma_S = 2.5$
$\Gamma_Y = 1.5$
$\Gamma_S = 2$
$\Gamma_Y = 2$
$\Gamma_S = 2$
$\Gamma_Y = 2.5$
$\Gamma_S = 2$
20
$A_Y$
0
$-1$
$B_Y$
1
$B_S = -0.5$
$B_S = 0$
$B_S = 0.5$
$A_S = 10$
$A_S = 5$
$A_S = 1$
$A_S = 0$

**Fig S7. Computer simulations using the new model over a wide range of combinations of seven parameters focused on the inductive effect (3)**

Computer simulations were performed using the new model under various settings of eight parameters, $51$ settings for $A_Y$ $(0 \leq A_Y \leq 20)$, $51$ settings for $B_Y$ $(-1 \leq B_Y \leq 1)$, $9$ settings for $\Gamma_Y$ and $\Gamma_S$ $((\Gamma_Y, \Gamma_S) = (1.5, 2)$, $(2, 2)$, $(2.5, 2)$, $(2, 2.5)$, $(2.5, 2.5)$, $(3, 2.5)$, $(2.5, 3)$, $(3, 3)$, or $(3, 3.5))$, $4$ settings for $A_S$ ($A_S = 0$, $1$, $5$, or $10$), and $3$ settings for $B_S$ ($B_S = 0$, $-0.5$, or $0.5$). $N$, $\alpha_Y$, and $\alpha_S$ were fixed to $1/3$, $2$, and $3$, respectively. The patterns obtained are displayed in the $A_Y B_Y$ space according to the color legend shown in Fig 2. Simulations were started by placing a single primordium.

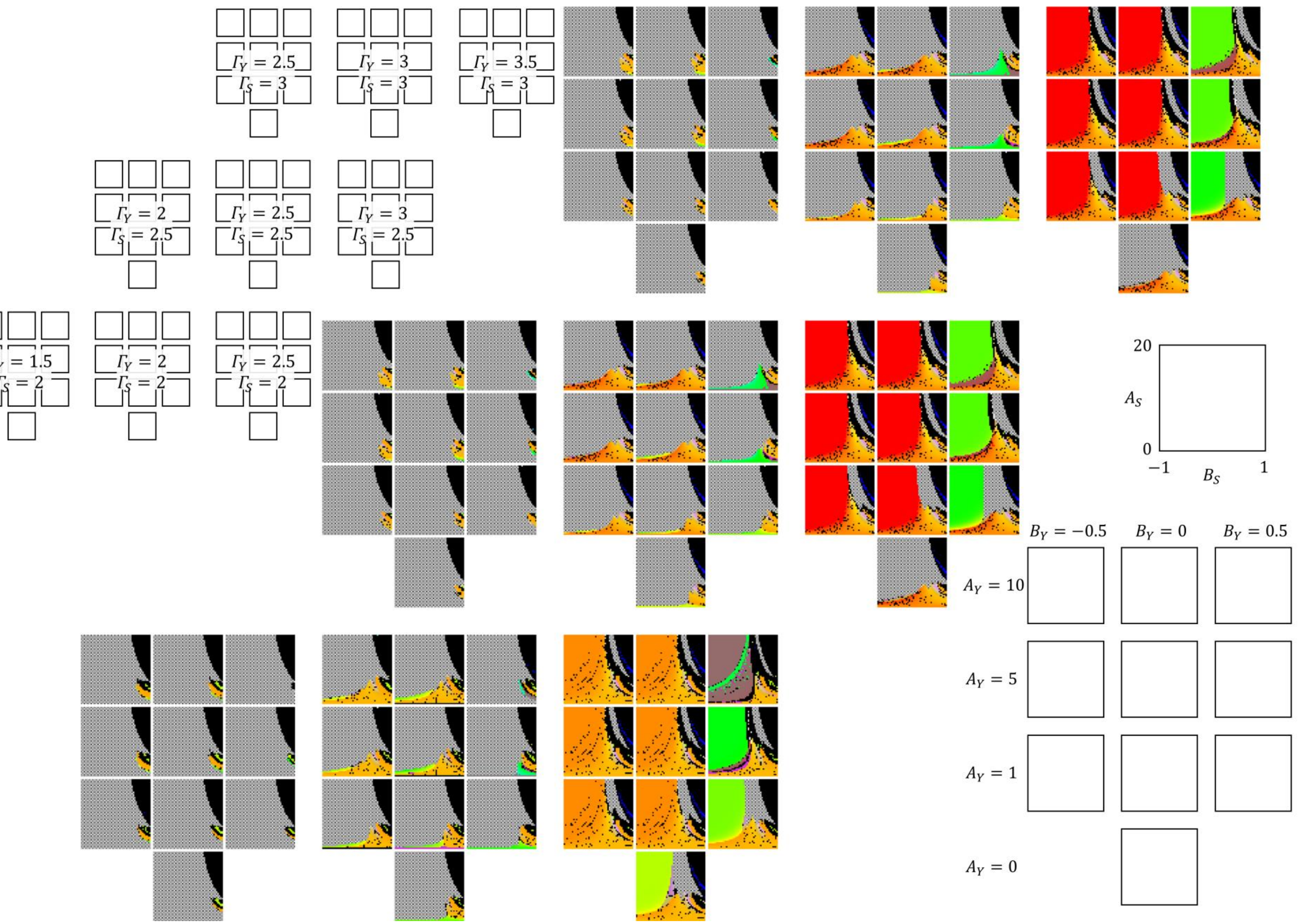

$\Gamma_Y = 2.5$ $\Gamma_S = 3$
$\Gamma_Y = 3$ $\Gamma_S = 3$
$\Gamma_Y = 3.5$ $\Gamma_S = 3$
$\Gamma_Y = 2$ $\Gamma_S = 2.5$
$\Gamma_Y = 2.5$ $\Gamma_S = 2.5$
$\Gamma_Y = 3$ $\Gamma_S = 2.5$
$\Gamma_Y = 2$ $\Gamma_S = 2$
$\Gamma_Y = 2.5$ $\Gamma_S = 2$
20
$A_S$
0
−1
$B_S$
1
$B_Y = -0.5$
$B_Y = 0$
$B_Y = 0.5$
$A_Y = 10$
$A_Y = 5$
$A_Y = 1$
$A_Y = 0$

**Fig S8. Computer simulations using the new model over a wide range of combinations of seven parameters focused on the inhibitory effect (1)**

Computer simulations were performed using the new model under various settings of eight parameters, $51$ settings for $A_S$ $(0 \leq A_S \leq 20)$, $51$ settings for $B_S$ $(-1 \leq B_S \leq 1)$, 9 settings for $\Gamma_Y$ and $\Gamma_S$ $((\Gamma_Y, \Gamma_S) = (1.5, 2)$, $(2, 2)$, $(2.5, 2)$, $(2, 2.5)$, $(2.5, 2.5)$, $(3, 2.5)$, $(2.5, 3)$, $(3, 3)$, or $(3, 3.5))$, $4$ settings for $A_Y$ ($A_Y = 0$, $1$, $5$, or $10$), and $3$ settings for $B_Y$ ($B_Y = 0$, $-0.5$, or $0.5$). $N$, $\alpha_Y$, and $\alpha_S$ were fixed to $1/3$, 4, and 3, respectively. The patterns obtained are displayed in the $A_S B_S$ space according to the color legend shown in Fig 2. Simulations were started by placing a single primordium.

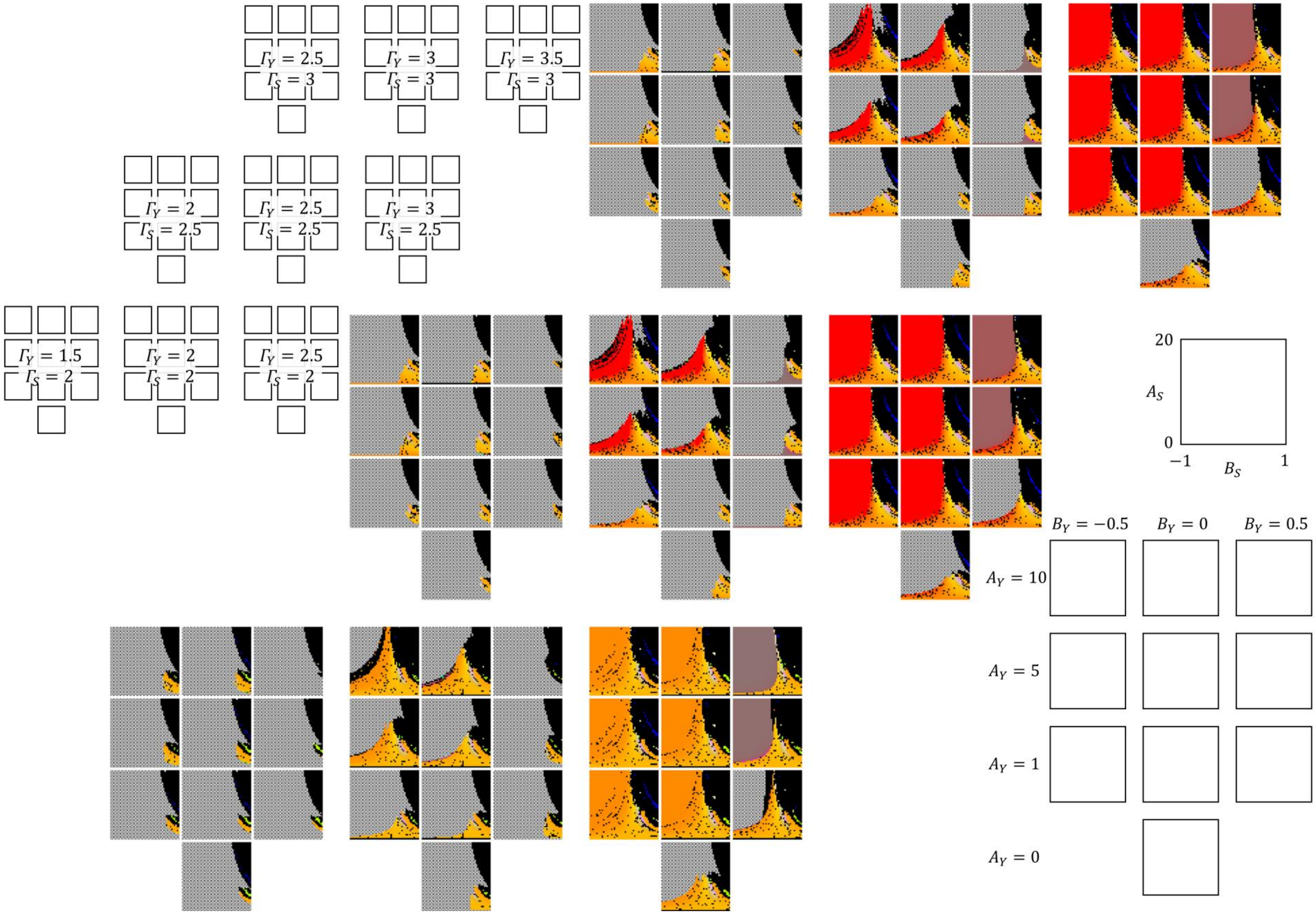

$\Gamma_Y = 2.5$ $\Gamma_S = 3$
$\Gamma_Y = 3$ $\Gamma_S = 3$
$\Gamma_Y = 3.5$ $\Gamma_S = 3$
$\Gamma_Y = 2$ $\Gamma_S = 2.5$
$\Gamma_Y = 2.5$ $\Gamma_S = 2.5$
$\Gamma_Y = 3$ $\Gamma_S = 2.5$
$\Gamma_Y = 1.5$ $\Gamma_S = 2$
$\Gamma_Y = 2$ $\Gamma_S = 2$
$\Gamma_Y = 2.5$ $\Gamma_S = 2$
20
$A_S$
0
−1
$B_S$
1
$B_Y = -0.5$
$B_Y = 0$
$B_Y = 0.5$
$A_Y = 10$
$A_Y = 5$
$A_Y = 1$
$A_Y = 0$

**Fig S9. Computer simulations using the new model over a wide range of combinations of seven parameters focused on the inhibitory effect (2)**

Computer simulations were performed using the new model under various settings of eight parameters, $51$ settings for $A_S$ $(0 \le A_S \le 20)$, $51$ settings for $B_S$ $(-1 \le B_S \le 1)$, 9 settings for $\Gamma_Y$ and $\Gamma_S$ $((\Gamma_Y, \Gamma_S) = (1.5, 2)$, $(2, 2)$, $(2.5, 2)$, $(2, 2.5)$, $(2.5, 2.5)$, $(3, 2.5)$, $(2.5, 3)$, $(3, 3)$, or $(3, 3.5))$, $4$ settings for $A_Y$ $(A_Y = 0,\ 1,\ 5$, or $10)$, and $3$ settings for $B_Y$ $(B_Y = 0,\ -0.5$, or $0.5)$. $N$, $\alpha_Y$, and $\alpha_S$ were fixed to $1/3$, 3, and 3, respectively. The patterns obtained are displayed in the $A_S B_S$ space according to the color legend shown in Fig 2. Simulations were started by placing a single primordium.

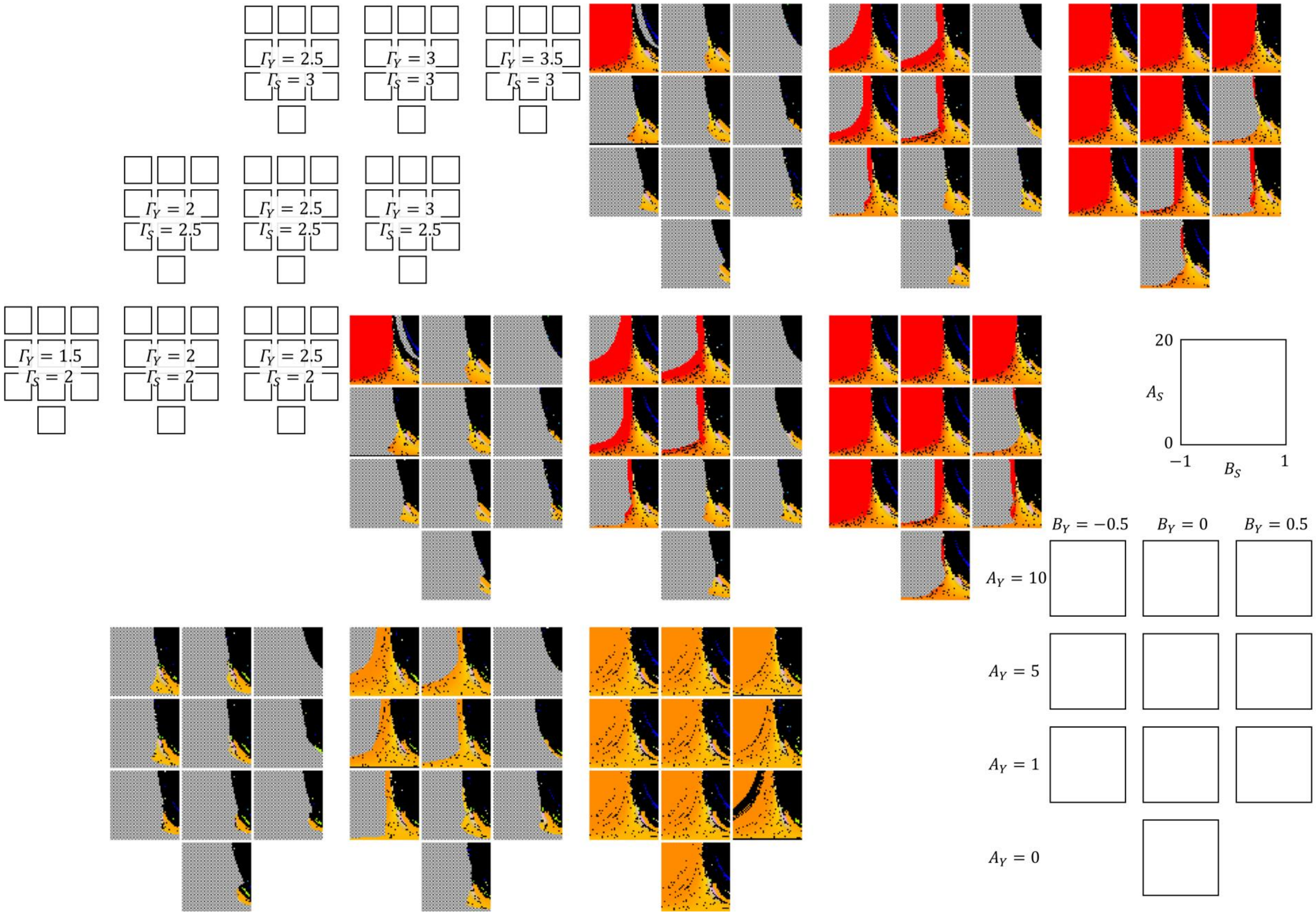

$\Gamma_Y = 2.5$ $\Gamma_S = 3$
$\Gamma_Y = 3$ $\Gamma_S = 3$
$\Gamma_Y = 3.5$ $\Gamma_S = 3$
$\Gamma_Y = 2$ $\Gamma_S = 2.5$
$\Gamma_Y = 2.5$ $\Gamma_S = 2.5$
$\Gamma_Y = 3$ $\Gamma_S = 2.5$
$\Gamma_Y = 1.5$ $\Gamma_S = 2$
$\Gamma_Y = 2$ $\Gamma_S = 2$
$\Gamma_Y = 2.5$ $\Gamma_S = 2$
20
$A_S$
0
−1
$B_S$
1
$B_Y = -0.5$
$B_Y = 0$
$B_Y = 0.5$
$A_Y = 10$
$A_Y = 5$
$A_Y = 1$
$A_Y = 0$

**Fig S10. Computer simulations using the new model over a wide range of combinations of seven parameters focused on the inhibitory effect (3)**

Computer simulations were performed using the new model under various settings of eight parameters, $51$ settings for $A_S$ ($0 \leq A_S \leq 20$), $51$ settings for $B_S$ ($-1 \leq B_S \leq 1$), 9 settings for $\Gamma_Y$ and $\Gamma_S$ ($(\Gamma_Y, \Gamma_S) = (1.5, 2)$, $(2, 2)$, $(2.5, 2)$, $(2, 2.5)$, $(2.5, 2.5)$, $(3, 2.5)$, $(2.5, 3)$, $(3, 3)$, or $(3, 3.5)$), $4$ settings for $A_Y$ ($A_Y = 0$, $1$, $5$, or $10$), and $3$ settings for $B_Y$ ($B_Y = 0$, $-0.5$, or $0.5$). $N$, $\alpha_Y$, and $\alpha_S$ were fixed to $1/3$, $2$, and $3$, respectively. The patterns obtained are displayed in the $A_S B_S$ space according to the color legend shown in Fig 2. Simulations were started by placing a single primordium.

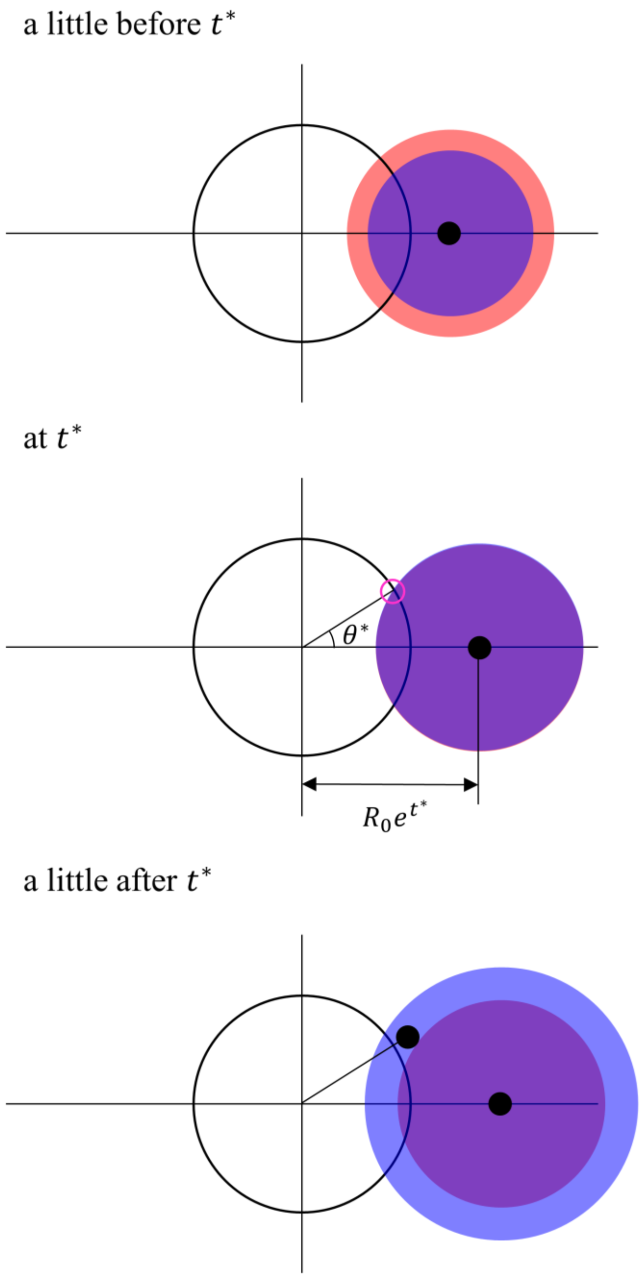


**Fig S11. Schematic illustration of the simplified situation of the new model**

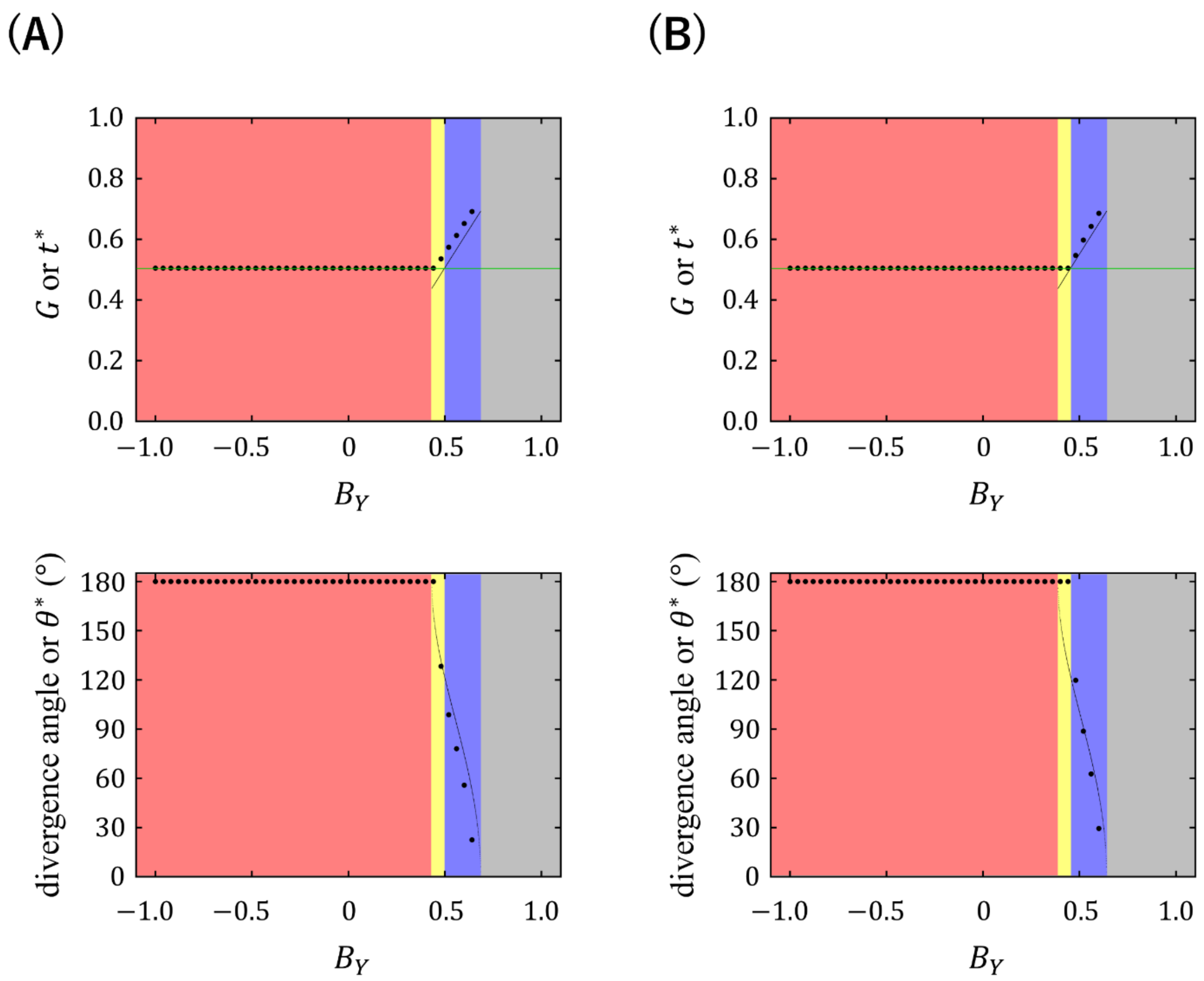


**Fig S12. Comparison between the solutions obtained from consideration of the simplified situation of the new model and the patterns generated by the computer simulations using the new model**

$t^*$ and $\theta^*$ were obtained for the simplified situation of the new model under various $B_Y$ values ($-1 \leq B_Y \leq 1$) with parameters $N = 1/3$, $\alpha_Y = 3$ or 4, $\alpha_S = 3$, $\Gamma_Y = 3.5$, $\Gamma_S = 3$, $A_Y = 20$, $A_S = 10$, and $B_S = 0$. Computer simulations using the new model were performed under the same parameter conditions. Large dots show $G$ and divergence angles of the phyllotactic patterns generated by the computer simulations, and small dots show the solutions for $t^*$ and $\theta^*$ if the solutions exist. Green lines indicate the value of $G_S$. Red zone: the range of $B_Y$ where the induction range always encompasses the inhibition range over $M$ and the solutions for $t^*$ and $\theta^*$ do not exist. Yellow zone: the range of $B_Y$ where the solutions for $t^*$ and $\theta^*$ exist and $t^*$ is smaller than $G_S$. Blue zone: the range of $B_Y$ where the solutions for $t^*$ and $\theta^*$ exist and $t^*$ is larger than $G_S$. Grey zone: the range of $B_Y$ where the induction range is always encompassed by the inhibition range over $M$ and the solutions for $t^*$ and $\theta^*$ do not exist. (A) $\alpha_Y$ was set to $4$. With this $\alpha_Y$ setting, costoid phyllotaxis can occur at some conditions in the simulations with the new model (left panel of Fig 9). (B) $\alpha_Y$ was set to $3$. With this $\alpha_Y$ setting, one-sided distichous phyllotaxis can occur at some conditions in the simulations with the new model (right panel of Fig 9).

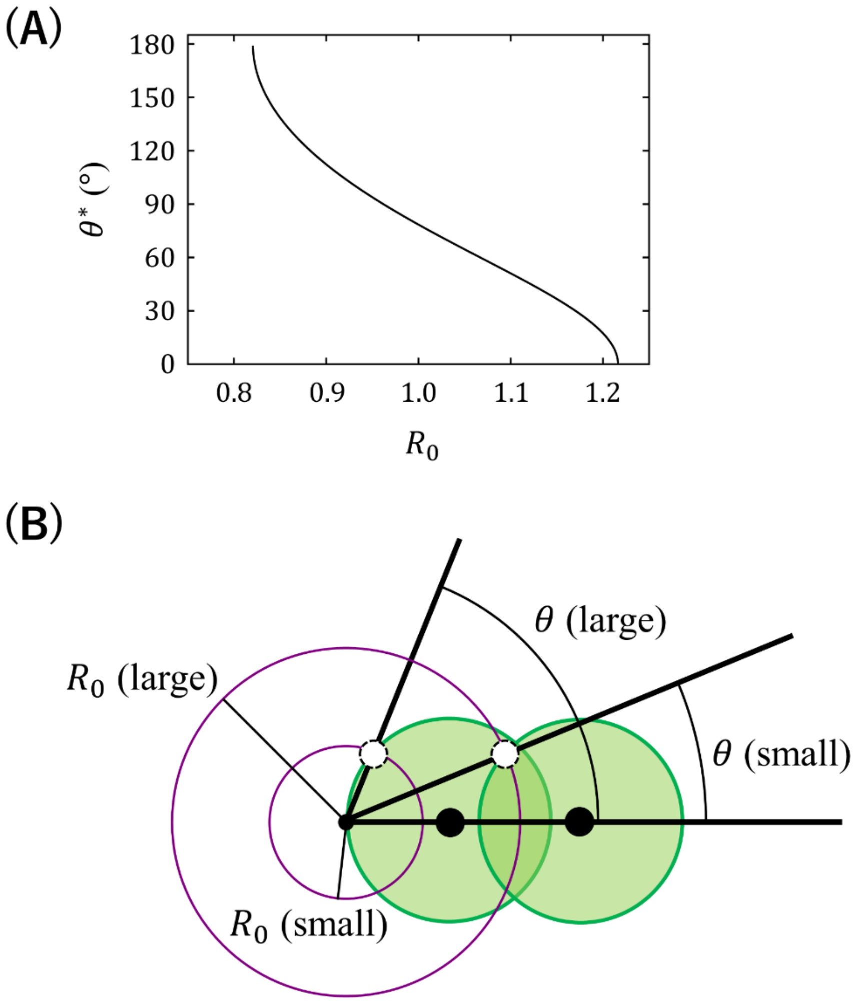


**Fig S13. Relationships between the SAM size and the divergence angle**

(A) Relationship between the SAM size $R_0$ and the predicted divergence angle $\theta^*$ in the simplified situation of the new model with parameters $N = 1/3$, $p = 3$, and $G = 0.6$. (B) Schematic illustration of the negative correlation between the SAM size and the divergence angle.
