## Supplementary material for "A new mathematical model of phyllotaxis to solve the genuine puzzle spiromonostichy": Text S1

**Text S1. Previous models.**

**EDC1**

The essential points of the EDC1 model are as follows (Yonekura et al., 2019).

1. The shoot apex is considered as a plane.
2. Each leaf primordium $L$ emits an inhibitory power, which generates an inhibitory field around it.
3. The inhibitory power increases as a function of the primordial age $t$.
4. The inhibitory field strength decreases as a function of the distance, $d$.
5. Formation of new primordia is restricted to the SAM periphery represented by the circle $M$ with the radius $R_0$ at the shoot apex.
6. New primordia are formed one by one at a regular time interval, $T$.
7. The point on $M$ at which the inhibitory field strength is smallest gives the radial position of the formation of a new primordium.
8. Primordia move away from the center of the shoot apex with a radial velocity of $V(r)$ that is proportional to the radial distance $r$ because of the exponential growth of the shoot apex.

At the time when the $n^{\text{th}}$ primordium $L_n$ is arising, for the position $(R_0 \cos\theta, R_0 \sin\theta)$ on the circle $M$, the inhibitory field strength $I(\theta)$ is calculated by summing the inhibitory effects from all preceding primordia $L_1$ to $L_{n-1}$ as follows:

$$I(\theta) \equiv \sum_{m=1}^{n-1} E\big(d_m(\theta)\big)F(t_m), \qquad (\mathrm{S1}-1)$$

where $d_m(\theta)$ is the distance from the position $(R_0 \cos\theta, R_0 \sin\theta)$ to the $m^{\text{th}}$ primordium $(r_m \cos\theta_m, r_m \sin\theta_m)$ and $t_m$ is the age of the $m^{\text{th}}$ primordium expressed as a value relative to $T$. $E\big(d_m(\theta)\big)$ and $F(t_m)$ give functions of the distance-dependent decrease of the inhibitory field strength and the age-dependent increase of the inhibitory power, respectively. These two functions are determined as:

$$E\big(d_m(\theta)\big) \equiv k\big(d_m(\theta)\big)^{-\eta} = k(R_0^2 + r_m^2 - 2R_0 r_m \cos(\theta - \theta_m))^{-\frac{\eta}{2}}, \qquad (\mathrm{S1}-2)$$

$$F(t_m) \equiv \frac{1}{1 + e^{-a(t_m - b)}}, \qquad (\mathrm{S1}-3)$$

where $k$, $\eta$, $a$, and $b$ are constants. Considering assumptions 5 and 7, the distance from the center of the shoot apex to the $m^{\text{th}}$ primordium ($r_m$) is expressed with the initial radial velocity $V_0$ as:

$$r_m = R_0 e^{\frac{V_0}{R_0}(n-m)T}. \qquad (\mathrm{S1}-4)$$

The total inhibitory field strength $I$ is expressed as:

$$I(\theta) = \frac{k}{R_0^{\eta}} \sum_{m=1}^{n-1} F(t_m)\{1 + e^{2(n-m)G} - 2e^{(n-m)G}\cos(\theta - \theta_m)\}^{-\frac{\eta}{2}}, \qquad (S1-5)$$

where $G$ is defined as $G \equiv V_0T/R_0 = \ln(r_{m+1}/r_m)$. Morphometrically, $r_{m+1}/r_m$ is identical to the "plastochron ratio" introduced by Richards (1951).

The point $(R_0 \cos\theta, R_0 \sin\theta)$ where $I(\theta)$ is smallest is chosen for the position of a new primordium. Note that $\eta$ and $G$ are the only relevant parameters that influence the behavior of $I(\theta)$ in DC1.

**EDC2**

The essential points of the EDC2 are as follows (Yonekura et al., 2019).

1. The shoot apex is considered as a cone with an apical angle of $\psi$.
2. Each leaf primordium $L$ emits an inhibitory power, which generates an inhibitory field around it.
3. The inhibitory power increases as a function of the primordial age $t$.
4. The inhibitory field strength decreases as a function of the distance, $d$.
5. Formation of new primordia is restricted to the SAM periphery represented by the circle $M$ with a distance of $R_0$ from the conical vertex.
6. When the inhibitory field strength falls below a given threshold $E_{th}$ somewhere on $M$, a new primordium is formed at that point.
7. Primordia move away from the center of the shoot apex with a radial velocity of $V(r)$ that is proportional to the radial distance $r$ because of the exponential growth of the shoot apex.

Positions on the conical surface are expressed in spherical coordinates $\left(r, \frac{\psi}{2}, \theta\right)$ (Fig 2-2B).

Because of assumption 7, the distance from the center of the shoot apex to the $m^{\text{th}}$ primordium on the conical surface ($r_m$) is expressed with the time after its emergence $T_m$ and the initial radial velocity $V_0$ as:

$$r_m = R_0 e^{\frac{V_0}{R_0}T_m}. \qquad (S1-6)$$

By using $t_m \equiv T_m V_0/R_0$, a standardized age of the $m^{\text{th}}$ primordium defined as the product of $T_m$ and the relative SAM growth rate $V_0/R_0$, $r_m$ is more simply expressed as:

$$r_m = R_0 e^{t_m}. \qquad (S1-7)$$

The inhibitory field strength $I(\theta)$ at the position $\left(R_0, \frac{\psi}{2}, \theta\right)$ on $M$ is calculated by summing the inhibitory effects from all preceding primordia $L_1$ to $L_{n-1}$ as follows:

$$I(\theta) \equiv \sum_{m=1}^{n-1} E\left(\frac{d_m(\theta)}{d_0}\right) F(t_m), \qquad (S1-8)$$

where $d_m$ is the distance from the $m^{\text{th}}$ primordium to the position $\left(R_0, \frac{\psi}{2}, \theta\right)$ and $d_0$ is the maximum distance within which an existing primordium excludes a new primordium. Two functions $E$ and $F$ are defined as:

$$E(x) \equiv E_{th} \frac{-1 + (\tanh \alpha x)^{-1}}{-1 + (\tanh \alpha)^{-1}}, \qquad (S1-9)$$

$$F(t_m) \equiv \frac{1}{1 + e^{-A(t_m - B)}}, \qquad (S1-10)$$

where $\alpha$, $A$, and $B$ are constants. If $I(\theta) < E_{th}$, a new primordium is placed at the position $\left(R_0, \frac{\psi}{2}, \theta\right)$. Throughout this study, $E_{th} = 1$.

The EDC2 model is characterized by five parameters: $\alpha$, $N \equiv \sin\frac{\psi}{2}$, $\Gamma \equiv \frac{d_0}{R_0\sqrt{N}}$, $A$, and $B$. These parameters represent the steepness of the decline of the inhibitory effect around the threshold, the flatness of the shoot apex, the ratio of the inhibition range to the SAM size, the steepness of the age-dependent increase of the inhibitory power, and the timing of age-dependent increase of inhibitory power, respectively.
